# Brain age estimation is a sensitive marker of processing speed in the early course of multiple sclerosis

**DOI:** 10.1101/651521

**Authors:** Einar A. Høgestøl, Gro O. Nygaard, Petter E. Emhjellen, Tobias Kaufmann, Mona K. Beyer, Piotr Sowa, Ole A. Andreassen, Elisabeth G. Celius, Nils Inge Landrø, Lars T. Westlye, Hanne F. Harbo

## Abstract

**Background and objectives:** Cognitive deficits in MS are common, also early in the disease course. We aimed to identify if estimated brain age from MRI could serve as an imaging marker for early cognitive symptoms in a longitudinal MS study.

**Methods:** A group of 76 MS patients (mean age 34 years, 71% females, 96% relapsing-remitting) was examined 1, 2 and 5 years after diagnosis. A machine-learning model using Freesurfer-processed T1-weighted brain MRI data from 3208 healthy controls, was applied to develop a prediction model for brain age. The difference between estimated and chronological brain age was calculated (brain age gap). Tests of memory, attention and executive functions were performed. Associations between brain age gap and cognitive performance were assessed using linear mixed effects (LME) models and corrected for multiple testing.

**Results:** LME models revealed a significant association between the Color Naming condition of Color-Word Interference Test and brain age gap (t=2.84, p=0.005).

**Conclusions:** In this study, decreased information processing speed correlated with increased brain age gap. Our findings suggest that brain age estimation using MRI provides a useful semi-automated approach applying machine learning for individual level brain phenotyping and correlates with information processing speed in the early course of MS.

## INTRODUCTION

MS is a chronic inflammatory disease of the CNS, mostly diagnosed in young adults. Cognitive deficits, affecting up to 70 % of all MS patients, are associated with psychiatric symptoms, reduced quality of life and ability to participate in work-related and social activities ^1^. Cognitive deficits may appear early in the disease course and are only mildly associated with physical disability ^2, 3^. The domains most frequently affected in MS are information processing speed and memory, followed by executive functions, verbal fluency and visuospatial processing ^4–6^.

MRI is an essential tool for diagnosing and monitoring of disease activity and progression in MS ^7, 8^. MRI is highly sensitive to MS related pathological processes such as inflammation, demyelination and loss of neurons and is an important tool for visualizing the neuropathological substrates in MS ^9^.

Cross-sectional studies have revealed cognitive deficits to be associated with several structural MRI (sMRI) markers. Longitudinal studies have shown correlation between corpus callosum atrophy ^10^, thalamus atrophy ^11, 12^, whole-brain atrophy ^13^ and reduction in grey and white matter volumes ^14, 15^ and cognitive performance. Especially, thalamus has shown to be highly susceptible to retrograde degeneration and scan-scan correlations with longitudinal data ^16^. Thalamus is recognized to be subject to atrophy from the earliest stages of MS, especially for primary progressive (PP) MS patients but also in general for MS patients ^14^. Thalamus structure has been linked to decline in cognitive performance in MS ^12^.

A multiparametric MRI study identified that deep reduced gray matter volume and regional white matter atrophy were the strongest predictors of overall cognitive dysfunction in MS ^17^. Cognitive decline is found to be associated with increase in T2 lesion volume, cerebral atrophy, microstructural damage and cortical lesions ^4^.

Despite extensive research efforts concerning MRI markers in MS, most studies show weak associations between neuroradiological disease markers and cognitive performance (the cognitive clinico-radiological paradox)^4, 18, 19^. There is a need to establish new MRI markers that robustly relate to cognitive function, with the ability to predict future progression and monitor the effects of treatments on the individual level ^4^.

Brain age estimation has emerged as a robust MRI marker, combining sensitive measures of MRI-based brain morphometry using machine learning models, to estimate an individual brain age when correlating with a large MRI data set of healthy controls ^20, 21^. Having an older-appearing brain is associated with advanced physiological and cognitive ageing and mortality in several neurodegenerative and neurodevelopmental disorders ^20, 22, 23^. To our knowledge, no studies elaborating cognitive function in association with brain age in MS have yet been performed.

By using data from our prospective, longitudinal study of newly diagnosed MS patients, we aimed to evaluate if regional brain age estimation is sensitive to subtle changes in cognitive performance.

## MATERIALS AND METHODS

### Participants

In total 76 MS patients were recruited as described previously ^15, 24^. The patients were diagnosed in 2009-2012 in accordance with the at that time current McDonald Criteria ^25^. The patients were examined at three time points after being diagnosed with MS; time point 1 after 15 months (±12, n=76), time point 2 after 28 months (±9, n=75) and time point 3 after 66 months (±13, n=62). No Evidence of Disease Activity (NEDA-3) was defined as absence of both clinical relapses, disease progression and new or contrast enhancing lesions on MRI. Disease-modifying treatments (DMTs) were categorized within the following groups; no treatment (group 0); glatiramer acetate, interferons, teriflunomide or dimetylfumarate (group 1) and fingolimod, natalizumab or alemtuzumab (group 2). The disease course was defined according to the Lublin criteria ^26^.

### MRI acquisition

All MS patients were scanned using the same 1.5 T scanner (Avanto, Siemens Medical Solutions; Erlangen, Germany) equipped with a 12-channel head coil for up to three times in the study with the same MRI scanning sequence across all timepoints. Structural MRI data were collected using a 3D T1-weighted MPRAGE (Magnetization Prepared Rapid Gradient Echo) sequence, with the following parameters: TR (repetition time) / TE (echo time) / flip angle / voxel size / FOV (field of view) / slices / scan time / matrix / time to inversion = 2400 ms / 3.61 ms / 8° / 1.20 × 1.25 × 1.25 mm / 240 / 160 sagittal slices / 7:42 minutes / 192 × 192 / 1000 ms.

### MRI pre and postprocessing

The MRI pre and postprocessing for this data has been described previously ^24^. Manual quality control of the MRI scans from patients was performed by trained research personnel to identify and edit segmentation errors where possible (done for 15 MRI scans) and exclude data of insufficient quality or missing sequences (11 MRI scans). Lesions have been shown to not substantially influence the output data from Freesurfer or the brain age estimations ^24, 27^. Detailed information on MRI variables is given in supplementary material.

### Brain age estimation model

Detailed information on the brain age estimation model has been described previously ^21, 24^. To summarize we utilized a training set based on MRI scans from 3208 HC (54 % women, mean age 47.5 (±19.8) years, age range 12-95 years) obtained from several publicly available datasets and processed in the same MRI pipeline (Supplementary Fig. 1). Based on robust performance in previous machine learning competitions we chose the xgboost package in R ^28^ to create the brain age estimation model. To estimate the performance of the estimation model, a 10-fold cross-validation showed consistent performance and generalizability for the combined model for both genders (r = 0.91, Supplementary Fig. 2).

### Neuropsychological assessment

The participants were evaluated using a battery of 13 neuropsychological tests at all three time points, detailed information regarding the tests can be found in the supplementary material. Results from baseline and the first follow-up have been published previously ^15, 29^. We used the raw scores in the analyses.

### Statistical analysis

We used R (R Core Team, Vienna, 2018, version 3.7.0) for statistical analyses. To assess reliability of brain age and cognitive tests across time we computed the intraclass correlation coefficient (ICC) using the R package “irr” (https://CRAN.R-project.org/package=irr). Figures were made using “ggplot2” ^30^ and “cowplot” (https://CRAN.R-project.org/package=cowplot) in R.

The linear mixed effects (LME) models were performed using the R package “nlme” (https://CRAN.R-project.org/package=nlme). All LME models accounted for age, gender and months since time point 1 ^31^. To control for multiple testing, we calculated the degree of independence between the resulting cognitive data only, by making a 13 × 13 correlation matrix based on the Pearson’s correlation between all pair-wise combinations of the cognitive variables. Utilizing the ratio of observed eigenvalue variance to its theoretical maximum, the estimated equivalent number of independent traits in our analyses was 9.0 ^32^. To control for multiple testing, our significance threshold was therefore adjusted accordingly from 0.05 to 5.6 × 10^−3 32^.

### Data availability

Summary data as published in this paper will be available, but other data are not publicly available because of patient privacy restrictions decided by the Regional Ethical Committee. We may apply for permission to share data with new collaborators, still adhering to patient privacy requirements of the “Law of Health Research”. All code needed to replicate our described analyses is available upon request from the corresponding author.

## RESULTS

### Participant demographics and characteristics

A full summary of the demographic and clinical characteristics is provided in Table 1. At time point 1, the MS patients had a mean age of 35.3 years (range 21.3-49.5 years), 71% were females and they were diagnosed with MS 15 months previously (range 0-36 months). Mean age increased to 36.3 (range 22.7-50.6 years) and 40.4 years (range 25.5-53.7 years) at time point 2 and 3, respectively. 96% were diagnosed with relapsing-remitting MS (RRMS). NEDA-3 was achieved for a subset of 53% and 44% at time point 2 and 3, respectively. The proportion of patients using group 1 DMTs decreased during follow-up period with 65%, 48 % and 37% at time point 1, 2 and 3. Correspondingly, the use of group 2 DMTs increased among the subjects in the same time period from 13% at time point 1 to 23% and 32% at time point 2 and 3.

**Table 1.** Demographic and clinical characteristics of the multiple sclerosis patients.

|  | Time point 1<br>*n* = 76 | Time point 2<br>*n* = 75 | Time point 3<br>*n* = 62 |
| --- | --- | --- | --- |
| **(a) Demographic and clinical characteristics** |  |  |  |
| Female, *n* (%) | 54 (71) | 54 (72) | 44 (71) |
| Age, mean years (range) | 35.3 (21-49) | 36.3 (22-50) | 40.5 (25-53) |
| EDSS, median (SD, range) | 2.0 (0.9, 0-6) | 2.0 (0.9, 0-4) | 2.0 (1.3, 0-6) |
| Number of total attacks, mean (SD, range) | 1.8 (1.0, 0-5) | 2.0 (1.0) | 2.6 (1.3) |
| Months since MS diagnosis, mean (SD, range) | 15.2 (9.7, 0-34) | 27.8 (9.1, 13-49) | 66.4 (13.4, 17-95) |
| Years since first symptom, mean years (SD, range) | 4.8 (4.4, 0.2-19.4) | 5.9 (4.3, 1.3-20.6) | 10.0 (4.6 (4.5-23.6) |
| NEDA-3, *n* (%) | - | 40 (53) | 27 (44) |

| Multiple sclerosis classification |  |  |  |
| --- | --- | --- | --- |
| RRMS, n (%) | 73 (96) | 72 (96) | 60 (96) |
| PPMS, n (%) | 2 (3) | 2 (3) | 1 (2) |
| SPMS, n (%) | 1 (1) | 1 (1) | 1 (2) |
| Education |  |  |  |
| Years, mean (SD, range) | 15.3 (2.3, 10-18) | - | 16.0 (1.9, 10-18) |
| > 15 years education, n (%) | 53 (70) | - | 50 (81) |
| DMT |  |  |  |
| Group 0, n (%) | 17 (22) | 22 (29) | 19 (31) |
| Group 1, n (%) | 49 (65) | 36 (48) | 23 (37) |
| Group 2, n (%) | 10 (13) | 17 (23) | 20 (32) |
| (b) Self-reported questionnaires |  |  |  |
| FSS, mean (SD) | 4.2 (1.7) | 3.8 (1.9) | 4.1 (1.9) |
| Clinically significant fatigue (FSS mean $\geq 4$), n (%) | 38 (51) | 25 (45) | 33 (53) |
| BDI-II sum, mean (SD) | 9.1 (6.6) | 8.2 (6.2) | 7.8 (6.0) |
| Clinically significant depressive symptoms (BDI sum $\geq 14$), n (%) | 23 (31) | 13 (24) | 11 (17) |
| BDI-II: Beck Depression Inventory-II, DMT: Disease Modifying Treatment, FSS: Fatigue Severity Scale, Group 0; No Treatment, Group 1; Glatiramer Acetate, Interferons, Teriflunomide or Dimethylfumarate, Group 2; Fingolimod, Natalizumab or Alemtuzumab, NEDA: No Evidence of Disease Activity, PPMS: Primary progressive MS, RRMS: Relapsing-remitting MS, SPMS: Secondary progressive MS |  |  |  |

### Changes in brain age and brain morphometry over time

Summary statistics from the brain morphometry, cognitive tests and corresponding principal component analysis (PCA) (see supplementary material) are provided in Table 2. Estimated brain age gap (BAG, the difference between estimated and chronological brain age). increased from 2.8 at time point 1 to 4.6 at time point 3 (LME estimated effect of time: t=0.36, p=0.72). Both thalamus volume (t=-2.27, p=0.03) and normalized thalamus volume (t=2.73, p=7.2 × 10^−3^) showed significant longitudinal decrease. We also found significant decrease in white matter volume (t=-2.30, p=0.02), normalized white matter volume (t=-3.56, p=5.3 × 10^−4^) and normalized brain volume (t=-2.60, p=0.01). Raw morphometric volumes and normalized volumes showed significant correlations with white matter volume (r=0.49, p=9.8 × 10^−13^) and thalamus (r=0.59, p=2.2 × 10^−16^) and non-significant correlations with total brain volume (r=0.08, p=0.25) and grey matter volume (r=0.07, p=0.37).

|  | Time point 1 | Time point 2 | Time point 3 | All time points (LME models) |  |  |
| --- | --- | --- | --- | --- | --- | --- |
|  | *n* = 76 | *n* = 75 | *n* = 62 | *t* | *p* | Annual change |
| (a) MRI characteristics |  |  |  |  |  |  |
| Intracranial volume, mean mL (SD) | 1593.3 (142.6) | 1593.3 (142.6) | 1593.3 (142.6) | - | - |  |
| Brain age, mean years (SD) | 38.2 (12.5) | 39.9 (13.0) | 45.1 (13.3) | 0.36 | 0.72 |  |
| BAG, mean (SD) | 2.8 (9.0) | 3.3 (9.4) | 4.6 (9.8) | 0.36 | 0.72 |  |
| Brain volume, mean mL (SD) | 1144.1 (94.7) | 1140.6 (98.9) | 1123.0 (98.0) | -1.59 | 0.11 |  |
| WM volume, mean mL (SD) | 526.4 (52.7) | 526.1 (54.1) | 519.3 (56.8) | -2.30 | 0.02 | -1.84 ml |
| GM volume, mean mL (SD) | 642.9 (49.4) | 639.8 (52.1) | 628.3 (51.1) | -0.58 | 0.56 |  |
| Thalamus volume, mean mL (SD) | 14.8 (1.5) | 14.7 (1.5) | 14.3 (1.6) | -2.27 | 0.03 | -0.06 ml |
| Normalized brain volume (SD) | 71.9 (2.5) | 71.7 (2.8) | 70.8 (2.6) | -2.60 | 0.01 | -0.001 |
| Normalized WM volume (SD) | 33.0 (1.5) | 33.0 (1.6) | 32.7 (1.7) | -3.56 | $5.3 \times 10^{-4}$ | -0.001 |
| Normalized GM volume (SD) | 40.4 (1.9) | 40.2 (2.0) | 39.7 (2.0) | -0.53 | 0.59 |  |
| Normalized thalamus volume (SD) | 9.3 (0.7) | 9.3 (0.8) | 9.1 (0.8) | -2.73 | $7.2 \times 10^{-3}$ | $-3.6 \times 10^{-5}$ |
| WM hypointensities, mean ml (SD) | 3.3 (3.2) | 3.06 (3.1) | 3.9 (4.5) | -0.61 | 0.54 |  |
| **(b) Cognitive characteristics** |  |  |  |  |  |  |
| **Processing speed and working memory** |  |  |  |  |  |  |
| SDMT, mean (SD) | 52.4 (9.6) | 53.8 (10.0) | 54.5 (10.9) | 3.51 | $6.5 \times 10^{-4}$ | 0.87 |
| PASAT 3", mean (SD) | 45.8 (10.6) | - | 48.0 (9.0) | 1.62 | 0.11 |  |
| D-KEFS CWIT: Color Naming, mean seconds (SD) | 29.7 (5.2) | 28.4 (4.8) | 27.4 (4.1) | -4.25 | $4.3 \times 10^{-5}$ | -0.50 seconds |
| D-KEFS CWIT: Word Reading, mean seconds (SD) | 22.8 (4.6) | 21.4 (3.7) | 21.0 (3.5) | -3.47 | $7.2 \times 10^{-4}$ | -0.34 seconds |
| **Verbal memory** |  |  |  |  |  |  |
| CVLT-II: Immediate free recall for list A, n (SD) | 61.6 (10.8)<sup>3</sup> | 65.0 (9.6)<sup>4</sup> | 64.9 (10.5)<sup>3</sup> | 2.46 | 0.02 | 0.73 |
| CVLT-II: Immediate free recall for list B, n (SD) | 7.4 (2.8)<sup>3</sup> | 9.3 (3.2)<sup>4</sup> | 8.0 (2.9)<sup>3</sup> | 1.85 | 0.07 |  |
| CVLT-II: Short-delay free recall for list A, n (SD) | 13.2 (3.2)<sup>3</sup> | 14.1 (2.6)<sup>4</sup> | 14.0 (2.5)<sup>3</sup> | 1.83 | 0.07 |  |
| CVLT-II: Short-delay cued recall for list A, n (SD) | 13.4 (2.6)<sup>3</sup> | 14.4 (2.1)<sup>4</sup> | 14.6 (1.9)<sup>3</sup> | 3.33 | $1.2 \times 10^{-3}$ | 0.24 |
| **Visuospatial memory** |  |  |  |  |  |  |
| BVMT-R: Total recall, mean (SD) | 28.2 (5.6)<sup>5</sup> | 29.2 (4.9)<sup>5</sup> | 29.7 (4.4)<sup>5</sup> | 2.99 | $3.4 \times 10^{-3}$ | 0.35 |
| **Executive functions** |  |  |  |  |  |  |
| D-KEFS CWIT: Inhibition, mean seconds (SD) | 49.8 (13.4) | 46.7 (11.1) | 45.8 (9.5) | -3.38 | $9.9 \times 10^{-4}$ | -1.14 seconds |
| D-KEFS CWIT: Inhibition/Switching, mean seconds (SD) | 59.4 (18.9) | 53.0 (11.1) | 55.4 (11.1) | -2.02 | 0.05 | -0.86 seconds |
| *Inhibition minus mean of Color Naming and Word Reading, mean seconds (SD)* | 23.5 (11.9) | 25.3 (12.5) | 21.6 (7.7) | -2.16 | 0.03 | -0.72 seconds |
| *Inhibition/Switching minus mean of Color Naming and Word Reading, mean seconds (SD)* | 33.1 (17.8) | 28.1 (9.3) | 31.2 (9.7) | -1.10 | 0.28 |  |
| COWAT: Letter fluency, total, mean (SD) | 44.4 (10.2)<sup>6</sup> | 39.9 (8.9)<sup>7</sup> | 43.4 (11.9)<sup>6</sup> | 1.03 | 0.30 |  |
| COWAT: Category fluency, total, mean (SD) | 23.8 (4.7)<sup>8</sup> | 21.9 (4.3)<sup>9</sup> | 24.7 (5.0)<sup>8</sup> | 2.63 | $9.7 \times 10^{-3}$ | 0.43 |
| **Principal Component analysis** |  |  |  |  |  |  |
| PC 1, mean (SD) | 0.67 (2.3) | -0.24 (1.9) | -0.43 (2.0) | -4.38 | $2.6 \times 10^{-5}$ | -0.23 |
| PC 2, mean (SD) | 0.00 (1.7) | -0.16 (1.2) | 0.04 (1.1) | 1.34 | 0.18 |  |
| Longitudinal changes are estimated using LME models, accounting for time since time point 1, age and gender. Annual change is reported where we found significant longitudinal change (adjusted for age and gender).<br>BAG: Brain age gap, BVMT-R: Brief Visuospatial Memory Test – Revised Edition, COWAT: Controlled Oral Word Association Test, CVLT-II: California Verbal Learning Test – Second Edition, D-KEFS CWIT: Delis-Kaplan Executive Function System Color-Word Interference Test, PASAT: Paced Auditory Serial Addition Test, SDMT: Symbol Digit Modalities Test.<br><sup>1</sup>Written version; <sup>2</sup>Oral version; <sup>3</sup>Standard form; <sup>4</sup>Alternate form; <sup>5</sup>BVMT-R: Form 1, 2, and 3 were used at their respective time points; <sup>6</sup>FAS; <sup>7</sup>BIL; <sup>8</sup>Animals; <sup>9</sup>Clothes |  |  |  |  |  |  |

### Longitudinal cognitive changes

LME models revealed significant longitudinal improvements in performance for the following cognitive tests: SDMT (t=3.51, p=6.5 × 10^−4^), immediate free recall list for list A of California Verbal Learning Test – Second Edition (CVLT-II) (t=2.46, p=0.02), short-delay cued recall trail for list A of CVLT-II (t=3.33, p=1.2 × 10^−3^), Brief Visuospatial Memory Test -Revised (BVMT-R) (t=2.99, p=3.4 × 10^−3^), category fluency test (t=2.63, p=9.7 × 10^−3^), the Color Naming (t=-4.25, p=4.3 × 10^−5^) (Fig. 1A; Fig. 1C), Word Reading (t=-3-47, p=7.2 × 10^−4^) (Fig. 1B; Fig. 1D), Inhibition (t=-3.38, p=9.9 × 10^−4^) and Inhibition/Switching (t=-2.02, p=0.05) conditions of the Color-Word Interference Test (CWIT). In addition, the difference between the Inhibition condition of CWIT and the mean of Color Naming and Word Reading conditions decreased over time (t=-2.16, p=0.03). The cognitive tests showed overall high ICC (0.32-0.97, Supplementary Table 2). When considering all three time points, the most reliable cognitive tests were SDMT (ICC=0.72, p=8.7 × 10^−22^), the Word Reading condition of CWIT (ICC=0.72, p=3.9 × 10^−22^) and the Color Naming condition of CWIT (ICC=0.66, p=1.1 × 10^−17^). Longitudinal changes for all the other cognitive tests are provided in Supplementary Fig. 6-12, while a correlation matrix for all the cognitive tests are provided in Supplementary Fig. 13.

**Figure 1.**
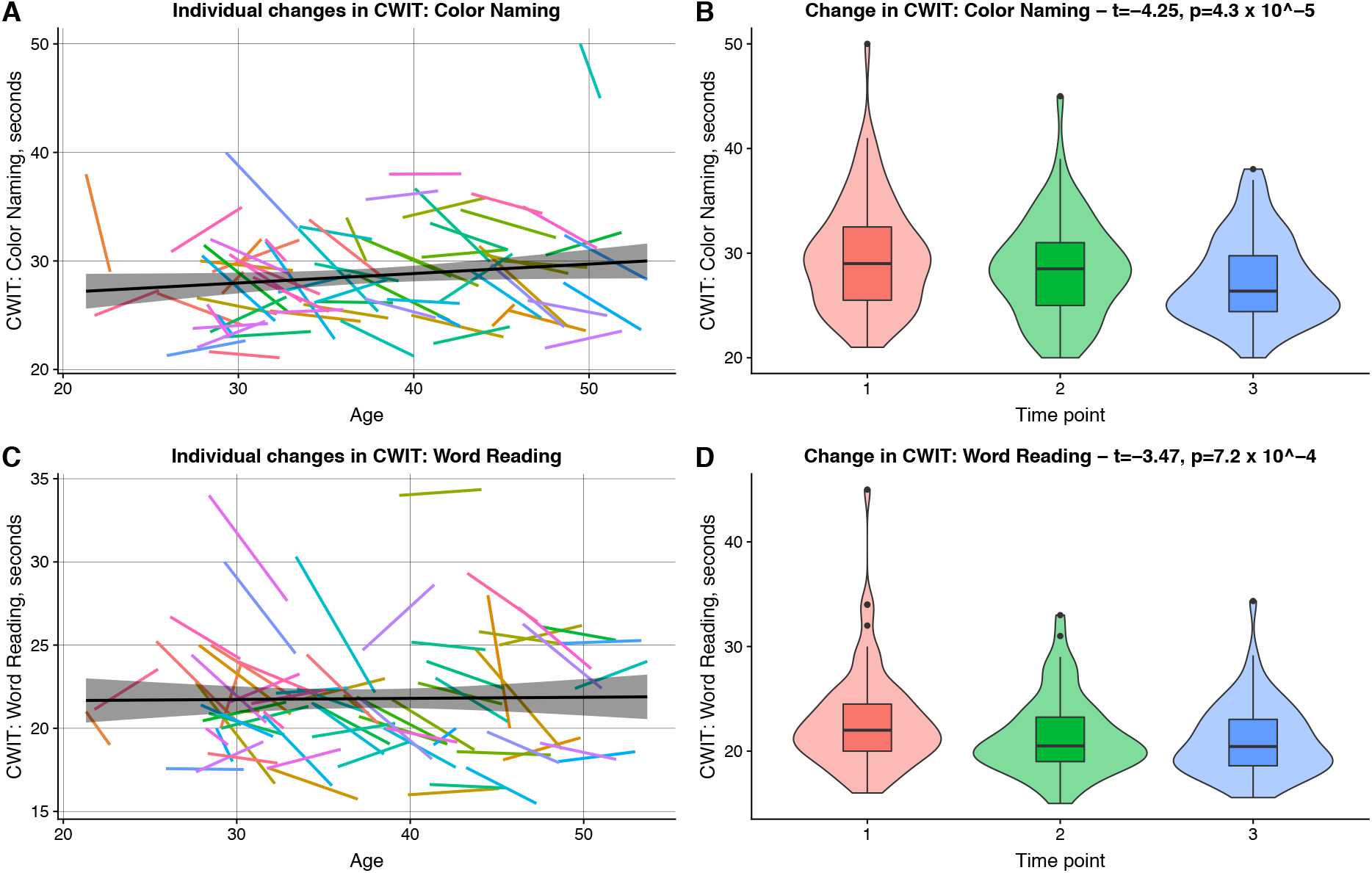
Longitudinal results for the Color Naming and Word Reading conditions of the Color-Word Interference Test. In (A) and (C) the individual regression lines for all subjects are depicted for the Color Naming and Word Reading conditions of CWIT, respectively. The black lines in (A) and (C) are the summarized regression lines for all data across all time points with the surrounding confidence interval. Furthermore, in (B) and (D) the boxplots for all subjects at all time points are shown for Color Naming and Word Reading, respectively. Both Color Naming (t=-4.25, p=4.3 × 10^−5^) and Word Reading (t=-3.47, p=7.2 × 10^−4^) displayed significantly decreasing results across the time points, as measured by LME models.

### Associations between brain age gap and cognitive performance

A summary of the multiple regressions analyses for associations between brain imaging markers and the cognitive tests are provided in Table 3 (see Supplementary Table 4 for complete results). After correcting for multiple testing there was a significant positive association between the Color Naming condition of CWIT and estimated BAG across all time points (t=2.84, p=5.4 × 10^−3^), indicating slower speed with higher brain age gap (Fig. 2). The associations between BAG and the Color Naming condition of CWIT remained after accounting for fatigue, years of education, disease duration, depressive symptoms, intracranial volume (ICV) and raw scores from the vocabulary task of Wechsler Abbreviated Scale of Intelligence. LME revealed no significant association between the longitudinal changes in the Color Naming condition of CWIT and BAG (stats for the interaction term: t=0.68, p=0.50).

**Figure 2.**
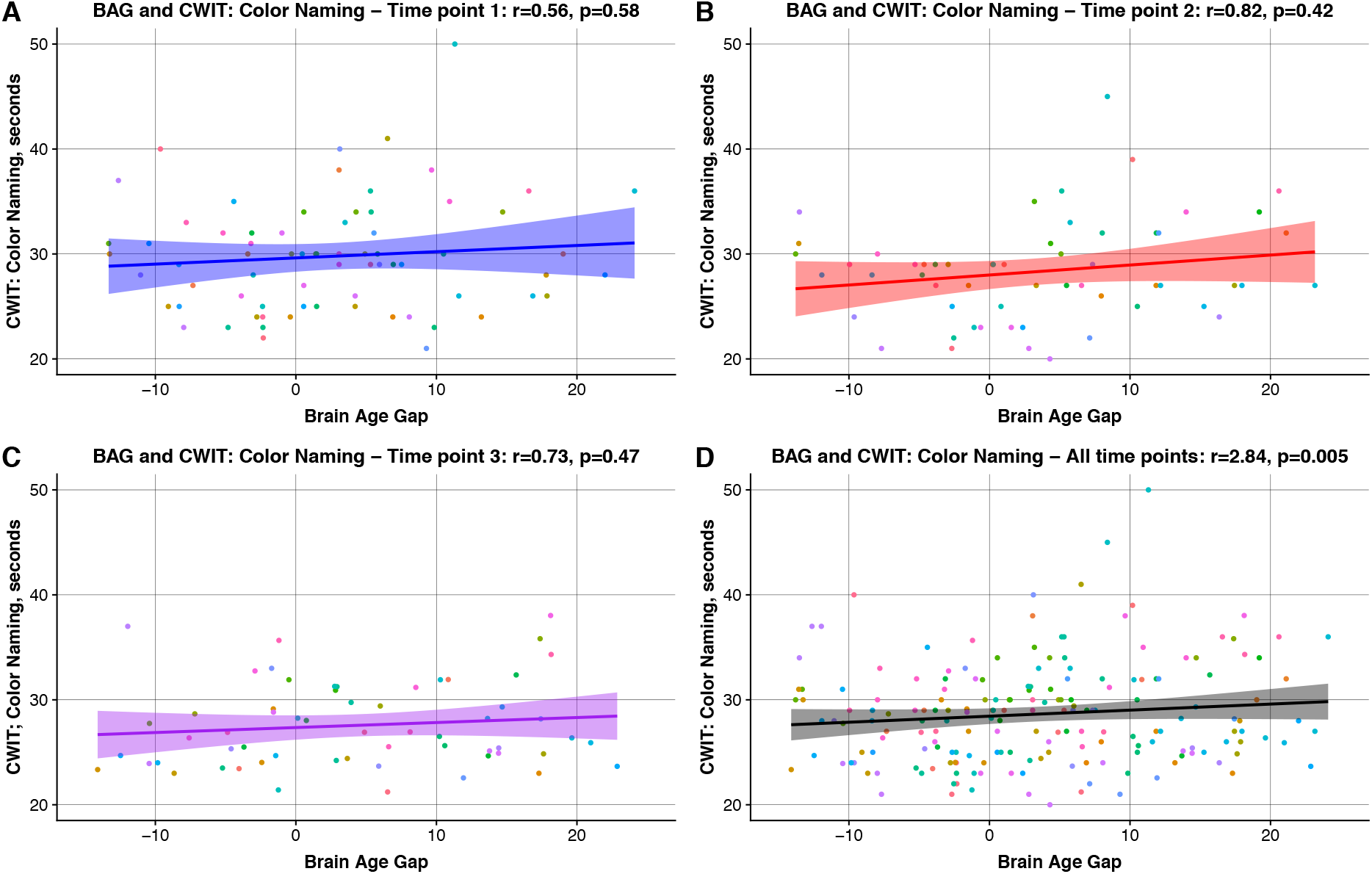
Associations between estimated brain age gap and the Color Naming condition of the Color-Word Interference Test. In (A), (B), and (C) the linear regression lines and the corresponding individual results for the Color Naming condition of CWIT across the estimated brain age gaps are shown at time point 1, 2 and 3, respectively. In (D) the summarized results for Color Naming at all time points across the estimated brain age gaps are displayed. Using an LME model there was a significant positive association between estimated BAG and Color Naming (t=2.84, p=5,4 × 10^−3^), also significant after correcting for multiple testing. The test results for all subjects are depicted using unique coloured circles for each subject.

**Table 3.** Summary of the correlations between the cognitive tests and MRI variables.

|  | MRI variables |  |  |  |
| --- | --- | --- | --- | --- |
|  | Brain age gap |  | Thalamus volume |  |
| Test variables | t | p | t | p |
| *Symbol Digit Modalities Test* |  |  |  |  |
| Time point 1 | -1.58 | 0.12 | 1.49 | 0.14 |
| Time point 2 | -0.48 | 0.63 | 0.67 | 0.51 |
| Time point 3 | -0.39 | 0.70 | 1.85 | 0.07 |
| All time points | -0.37 | 0.71 | 1.73 | 0.09 |
| *CVLT-II: Immediate free recall for list A* |  |  |  |  |
| Time point 1 | 0.34 | 0.73 | 0.99 | 0.33 |
| Time point 2 | 1.11 | 0.27 | 0.49 | 0.63 |
| Time point 3 | -0.27 | 0.79 | **3.1** | **$2.8 \times 10^{-3}$** |
| All time points | 0.23 | 0.82 | 1.94 | 0.05 |
| *CVLT-II: Immediate free recall for list B* |  |  |  |  |
| Time point 1 | -1.12 | 0.27 | **3.04** | **$3.4 \times 10^{-3}$** |
| Time point 2 | 1.05 | 0.30 | 0.87 | 0.39 |
| Time point 3 | 0.64 | 0.53 | 2.69 | 0.01 |
| All time points | 0.34 | 0.73 | 2.71 | $7.8 \times 10^{-3}$ |
| *CVLT-II: Short-delay free recall for list A* |  |  |  |  |
| Time point 1 | 0.58 | 0.56 | 0.77 | 0.45 |
| Time point 2 | 1.79 | 0.08 | -0.80 | 0.43 |
| Time point 3 | -0.10 | 0.92 | **2.98** | **$4.3 \times 10^{-3}$** |
| All time points | 1.11 | 0.27 | 0.99 | 0.32 |
| *CVLT-II: Short-delay cued recall for list A* |  |  |  |  |
| Time point 1 | 0.64 | 0.53 | -0.28 | 0.78 |
| Time point 2 | 1.17 | 0.25 | 0.86 | 0.39 |
| Time point 3 | -0.26 | 0.80 | **2.93** | **$4.9 \times 10^{-3}$** |
| All time points | 0.99 | 0.33 | 0.97 | 0.34 |
| *D-KEFS CWIT: Color naming* |  |  |  |  |
| Time point 1 | 0.56 | 0.58 | -1.26 | 0.21 |
| Time point 2 | 0.82 | 0.42 | 0.07 | 0.95 |
| Time point 3 | 0.73 | 0.47 | -2.16 | 0.036 |
| All time points | **2.84** | **$5.4 \times 10^{-3}$** | -1.75 | 0.08 |
| *D-KEFS CWIT: Word reading* |  |  |  |  |
| Time point 1 | 1.36 | 0.18 | **-3.28** | **$1.6 \times 10^{-3}$** |
| Time point 2 | 1.66 | 0.10 | -0.79 | 0.43 |
| Time point 3 | 0.79 | 0.43 | **-3.02** | **$3.8 \times 10^{-3}$** |
| All time points | 1.36 | 0.18 | **-3.09** | **$2.5 \times 10^{-3}$** |
| *First PCA component* |  |  |  |  |
| Time point 1 | 1.08 | 0.28 | -2.08 | 0.04 |
| Time point 2 | -1.02 | 0.31 | -1.02 | 0.31 |
| Time point 3 | 0.08 | 0.94 | **-3.80** | **$3.6 \times 10^{-4}$** |
| All time points | -0.42 | 0.68 | -2.23 | 0.03 |
| *Second PCA component* |  |  |  |  |
| Time point 1 | -1.62 | 0.11 | 1.43 | 0.16 |
| Time point 2 | -1.79 | 0.08 | 0.40 | 0.69 |
| Time point 3 | -0.33 | 0.75 | 0.35 | 0.73 |
| All time points | -1.54 | 0.13 | 1.28 | 0.20 |
| All longitudinal tests were run in R using the "nlme" package to calculate LME models, also accounting for months since time point 1, age and gender as fixed parameters. The linear models are calculated using the "stats" package in R, also |  |  |  |  |

accounting age and gender as fixed parameters. Brain age gaps are corrected for age and gender effects. Significant associations are marked as italic, while the associations still significant after correcting for multiple comparisons are marked bold and with thicker borders around.

### Associations between thalamus volume and cognitive performance

After correcting for multiple testing, LME revealed a significant negative association between the Word Reading condition of CWIT and thalamus volume across all time points (−3.09, p=2.5 × 10^−3^), suggesting slower performance with smaller thalamus volumes (Fig. 3). We found no significant association between the longitudinal changes in the Word Reading condition of CWIT and thalamus volume (stats for the interaction term: t=1.55, p=0.12). The association between thalamus volume and the Word Reading condition of CWIT remained after accounting for fatigue, years of education, disease duration, depressive symptoms, ICV or raw scores from the vocabulary task of Wechsler Abbreviated Scale of Intelligence.

**Figure 3.**
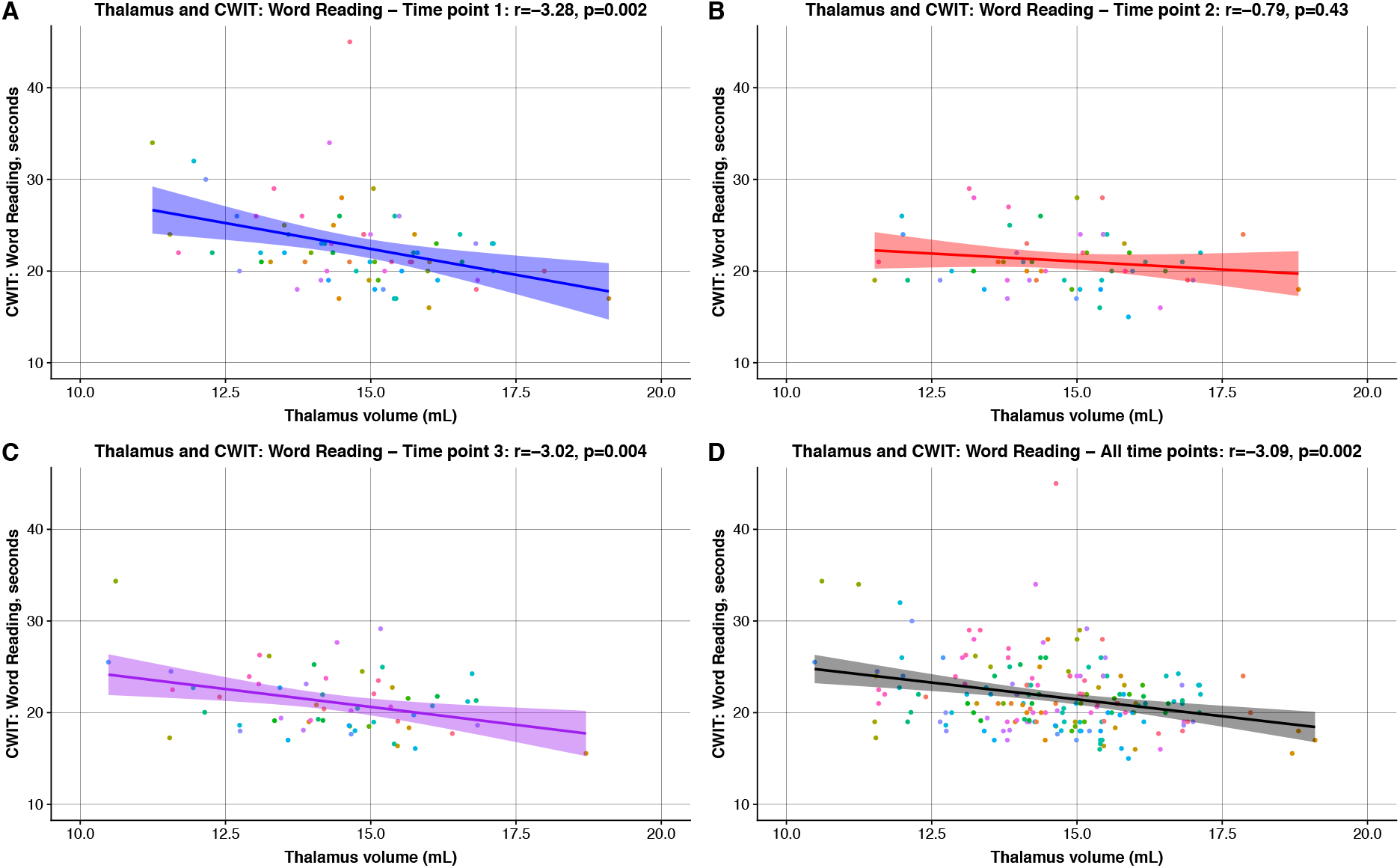
Associations between thalamus volume and the Word Reading condition of the Color-Word Interference Test. In (A), (B), and (C) the linear regression lines and the corresponding individual results for the Word Reading condition of CWIT across the estimated brain age gaps are shown at time point 1, 2 and 3, respectively. In (D) the summarized results for Word Reading at all time points across the estimated brain age gaps are displayed. Using an LME model there was a significant negative association between thalamus volume and Word Reading (t=-3.09, p=2.5 × 10^−3^), also significant after correcting for multiple testing. The test results for all subjects are depicted using unique coloured circles for each subject.

## DISCUSSION

Using established machine learning methods for brain age estimation, we tested if estimated brain age was associated with early cognitive decline in a longitudinal study of MS patients. In addition, we investigated associations with established MRI features and cognitive performance. We found that reduced information processing speed was associated with increased brain age gap and smaller thalamus volumes for patients in the early course of MS. Previous studies have shown that the first symptom of cognitive decline MS is impairment in information processing speed ^4, 33^, as our results also suggest with associations between MRI markers and decreased information processing speed. We show that brain age estimation correlates with cognitive decline measured by information processing speed in the early course of MS.

In our study, increasing brain age gap were significantly associated with slower performance for the Color Naming condition of CWIT across all time points, corresponding to an estimated 0.13 seconds increase in completion time for every year increase in brain age gap. This finding is in line with a recent study reporting lower processing speed as measured using the Stroop test with higher brain age gap in healthy individuals covering large parts of the adult lifespan ^22^. Supporting the sensitivity to individual differences in relevant clinical and cognitive traits, brain age gap has also been linked to negative symptoms in patients with schizophrenia, cognitive impairment in patients with dementia, and symptom burden in patients with MS ^21^. A study investigating brain age prediction over the lifespan revealed that individuals with major objective cognitive impairment had 2.1 years higher estimated brain age compared with a group of individuals with no objective cognitive impairment ^23^. Reduced information processing speed is known to appear early and with an increasing prevalence throughout the disease course of MS ^4^.

We found that reduced Word Reading condition of CWIT performance was significantly associated with smaller thalamus volume across time points corresponding to 0.87 seconds slower completion time for every ml reduction in thalamus volume. Previous studies have documented the importance of thalamic changes for cognitive performance in MS ^12^ while a longitudinal study has shown thalamic atrophy to be evident from the earliest stages in MS ^14^. Although the current results suggest a relevant role of the thalamus in processing speed, the extended brain networks supporting cognitive function are distributed and comprise large parts of the brain. Hence, combining information from various structures and imaging modalities is likely to improve sensitivity. Indeed, increased sensitivity and specificity to identify MS patients with severe cognitive impairment have been found when including other MRI modalities, such as resting-state functional MRI and diffusion tensor imaging ^12^, which is also in line with a study comparing the sensitivity to cognitive performance between various brain age estimations based on different combinations of structural MRI and diffusion tensor imaging ^22^.

Our results did not uncover relevant associations between imaging variables and SDMT. The SDMT is currently a widely used screening test for reduced information processing speed in a clinical setting and is the suggested sentinel test for cognitive impairment in MS ^6, 33^. The correlations between SDMT and the Color Naming and Word Reading conditions of CWIT were significant in our study. The lack of significant correlations between SDMT and the MRI markers across all time points might be due to the fact that our patients were at an early stage in their disease course. In accordance with the national guidelines for MS care in Norway, a large proportion of the MS patients in this study were treated with high efficacy DMTs from early on, possibly resulting in the higher number of NEDA patients in the study. As described earlier, on average our MS patients even performed better on some cognitive domains compared to healthy controls ^15^. In the same study, patients with NEDA had on average stable cognitive performances one year after the first visit ^15^. Findings from another MS sample also related lower depressive symptoms with improved cognitive performance ^3^.

No significant longitudinal correlations between changes in MRI parameters and cognitive test performance after the five year follow-up were found. One contributing factor to this might be the subtle cognitive and morphological changes observed in our patients. Significant improvements were found across the larger part of our administered cognitive tests at time point 1, 2 and 3. These improvements were most likely due to practice-related effects ^34^. Similar increases in test performance was also observed for SDMT in a study of Danish MS patients, where they found continuous enhancement after repeated monthly testing and more pronounced practice effect at lower EDSS levels ^35^.

Some additional limitations have to be considered when interpreting our results. First, we did not have access to longitudinal matched healthy controls, which would have enabled us to directly compare both the brain imaging and cognitive data across time points. Secondly, our current brain age estimation model was based solely on structural brain imaging data whereas some studies have shown increased precision when incorporating additional imaging modalities such as diffusion tensor imaging and functional MRI. In addition, the study does not allow us to make causal interference.

To conclude, this longitudinal MS study showed reduced information processing speed to be associated with increased brain age gap for patients in the early course of MS. Our results fit with previous findings, where reduced information speed is found to be an early symptom of cognitive decline in MS. In conclusion, we show that brain age estimation using MRI provides a useful semi-automated method for individual level brain phenotyping and correlates with cognitive decline measured by information processing speed.

## ACKNOWLEDGMENTS

We thank all patients participating in our studies. We acknowledge the collaboration with members of the Multiple Sclerosis Research Group and NORMENT at the University of Oslo and Oslo University hospital, especially Marte Wendel-Haga for her contribution at time point 2. We also thank the research assistants Kristin Liltved Grønsberg, May-Britt Gjengstø Utheim, Julia Timofeeva, Hedda Maurud and Siren Tønnessen who all contributed in the cognitive testing of the patients.

## AUTHOR CONTRIBUTIONS STATEMENT

EH, PE, GN, EC, NL, LW and HH contributed to the conception and design of the study. EH, PE, GN, TK, NC, PS, NL, LW and HH contributed to the acquisition and analysis of data. EH, PE, GN, NL, LW and HH drafted the text and figures. All authors contributed to the review and editing.

## SUPPLEMENTARY MATERIAL

Supplementary material is included in the supplemental files.

## ETHICS STATEMENT

This study was carried out in accordance with the recommendations of the Regional Committee for Medical and Health Research Ethics with written informed consent from all subjects. All subjects gave written informed consent in accordance with the Declaration of Helsinki. The protocol was approved by the South East Regional Committee for Medical and Health Research Ethics.

