## Supplementary Material for "Brain age estimation is a sensitive marker of processing speed in the early course of multiple sclerosis"

### MRI postprocessing

MRI variables were acquired from the Freesurfer output stats. We used the variable "Brain Segmentation Volume Without Ventricles from Surf" as measure of total brain volume, which excludes the brainstem. "Total grey matter volume" was used as an estimation of the grey matter (GM) volume. For the white matter (WM) volume we summarized the variables "brainstem", "cerebral white matter volume", "cerebellar white matter volume" and "corpus callosum". To account for individual differences, we also calculated normalized MRI variables by dividing the features by the "Estimated Total Intracranial Volume" provided by the Freesurfer output. We also included the WM hypointensities provided by Freesurfer.

### Neuropsychological assessments

The following cognitive domains were evaluated: Information processing speed and working memory were assessed using the Symbol Digit Modalities Test (SDMT) [1], with the total number of correctly substituted symbols in 90 seconds, the Paced Auditory Serial Addition Test (PASAT) – 3 seconds version [2]), with the total number of correctly performed calculations, and the Color-Word Interference Test (CWIT) subtest from the Delis-Kaplan Executive Function System (D-KEFS) [3]), with time to completion of the Color Naming and Word Reading conditions. Verbal memory was evaluated using the Norwegian version of the California Verbal Learning Test – Second Edition (CVLT-II) [4, 5], with the total number of correctly recalled words in each condition.

Visuospatial memory was evaluated using the Brief Visuospatial Memory Test - Revised (BVMT-R) [6], with the total number of points earned in the three immediate recall conditions. Executive functions were measured using the D-KEFS CWIT, with time to completion of the Inhibition and Inhibition/Switching conditions. Verbal fluency was assessed using the Controlled Oral Word Association Test (COWAT) [7]).

### Principal component analysis

The principal component analysis (PCA) was performed using the R package "factoextra" (https://cran.r-project.org/package=factoextra) and the embedded R package "stats". We performed a PCA of data from all 13 cognitive tests for all participants, imputing all missing data with the median value for that specific test at the specific time point. We extracted the subject scores for the two highest ranked PCA components for further analysis to test for associations with the MRI derived features (Supplementary Fig. 3, 4 and 5; Supplementary Table 1).

The first (PCA1, eigenvalue=4.60, variance explained=35.4%) and second (PCA2, eigenvalue=1.8, variance explained=14.1%) components of the PCA cumulatively explained 49.5% of the total variance in the cognitive data (Supplementary Fig. 3; Supplementary Table 1). Detailed information concerning the contributions of the cognitive tests in PCA1 and PCA2 are illustrated in Supplementary Fig. 3. 4. and 5. ICC for PCA1 (ICC=0.65, $p=6.6 \times 10^{-23}$) and PCA2 (ICC=0.62, $p=1.1 \times 10^{-20}$) showed high longitudinal reliability (Supplementary Table 3).

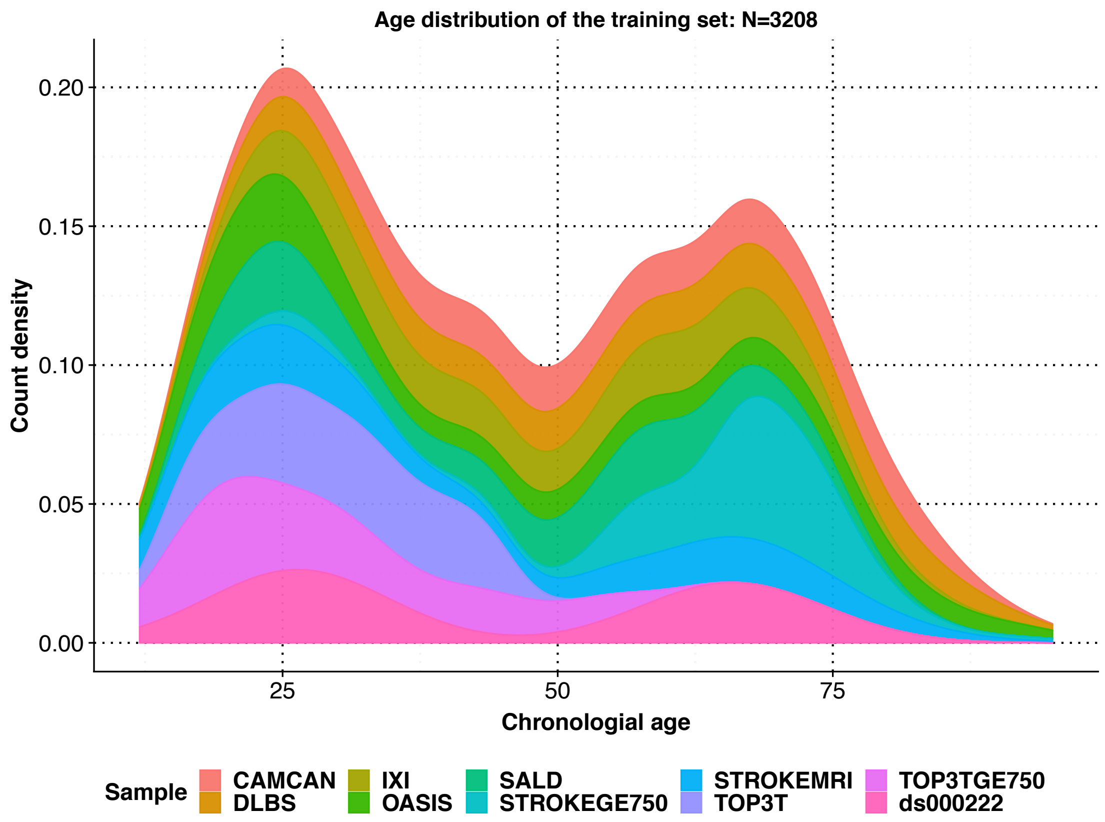


**Supplementary figure 1. Overview age distribution of the training set.** Histogram showing the age distribution of the training set, including the age distributions of the different cohorts using different colours. Information concerning the different sample cohorts (CAMCAN, DLBS, IXI, OASIS, SALD, STROKEGE750, STROKEMRI, TOP3T, TOP3TGE750 and ds000222) contributing to the training set are available in the article by Kaufman et al [8].

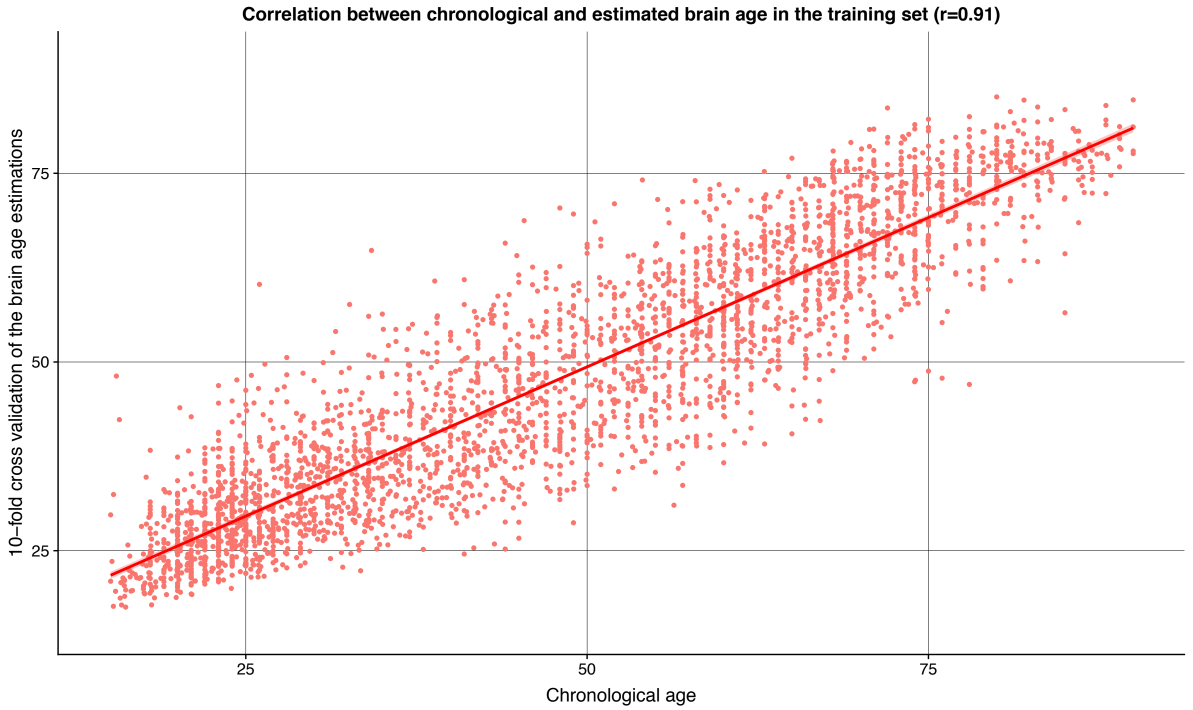


**Supplementary figure 2. Result of the brain age estimates from the training set.** Correlations between brain age estimations and age after performing a 10-fold cross validation in the training set (r = 0.91). We found a shift in the estimations <50 and >50 years of age, with increasing overestimation of age <50 years and increasing underestimation of age >50 years, which is a known effect when utilizing a machine learning model to estimate brain age [8, 9].

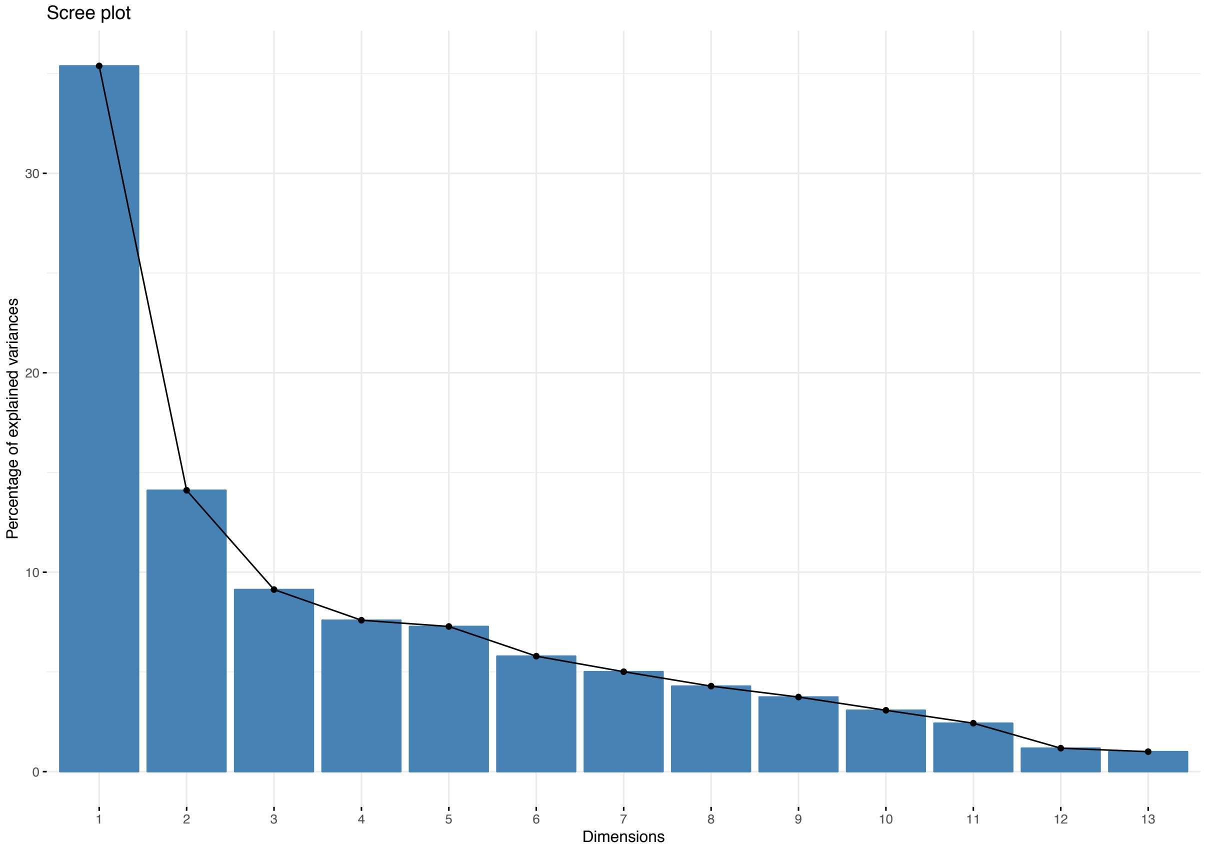


**Supplementary figure 3. Histogram showing all the PCA components and their explained percentage of explained variance in the data.** For detailed information see Supplementary table 4.

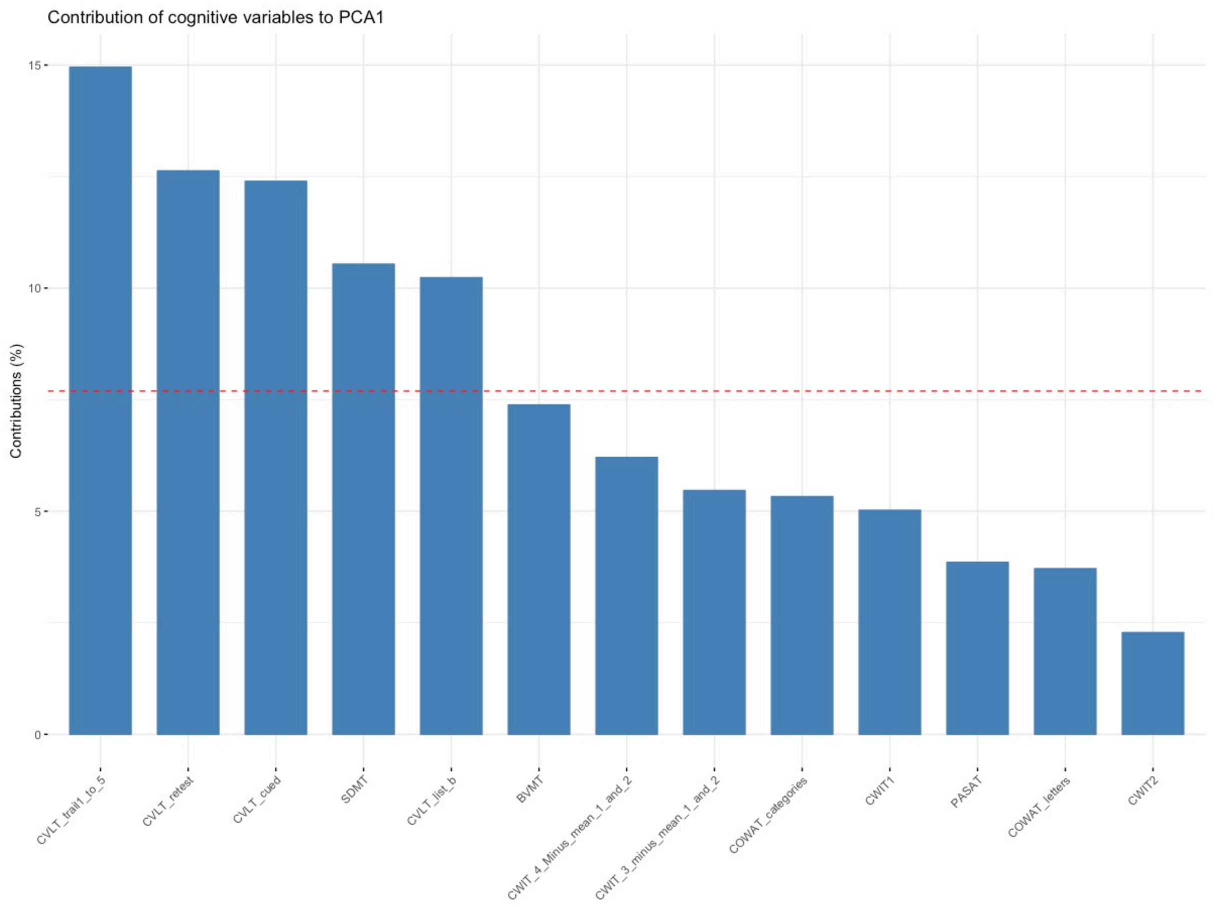


**Supplementary figure 4. Histogram visualizing the individual contributions of the cognitive variables in the first principal component.** CVLT_trail_1_to_5; CVLT-II: Immediate free recall for list A, CVLT_retest; CVLT-II: Short-delay free recall for list A, CVLT_cued; CVLT-II: Short-delay cued recall for list A, SDMT; Symbol Digit Modalities Test, CVLT_list_b; CVLT-II: Immediate free recall for list B, BVMT; Brief Visuospatial Memory Test – Revised Edition, CWIT_4_Minus_mean_1_and_2; Inhibition/Switching minus mean of Color Naming and Word Reading, CWIT_3_Minus_mean_1_and_2; Inhibition minus mean of Color Naming and Word Reading, COWAT_categories; COWAT: Category fluency, CWIT1; D-KEFS CWIT: Color Naming, PASAT; Paced Auditory Serial Addition Test, COWAT_letters; COWAT: Letter fluency, CWIT2; D-KEFS CWIT: Word Reading.

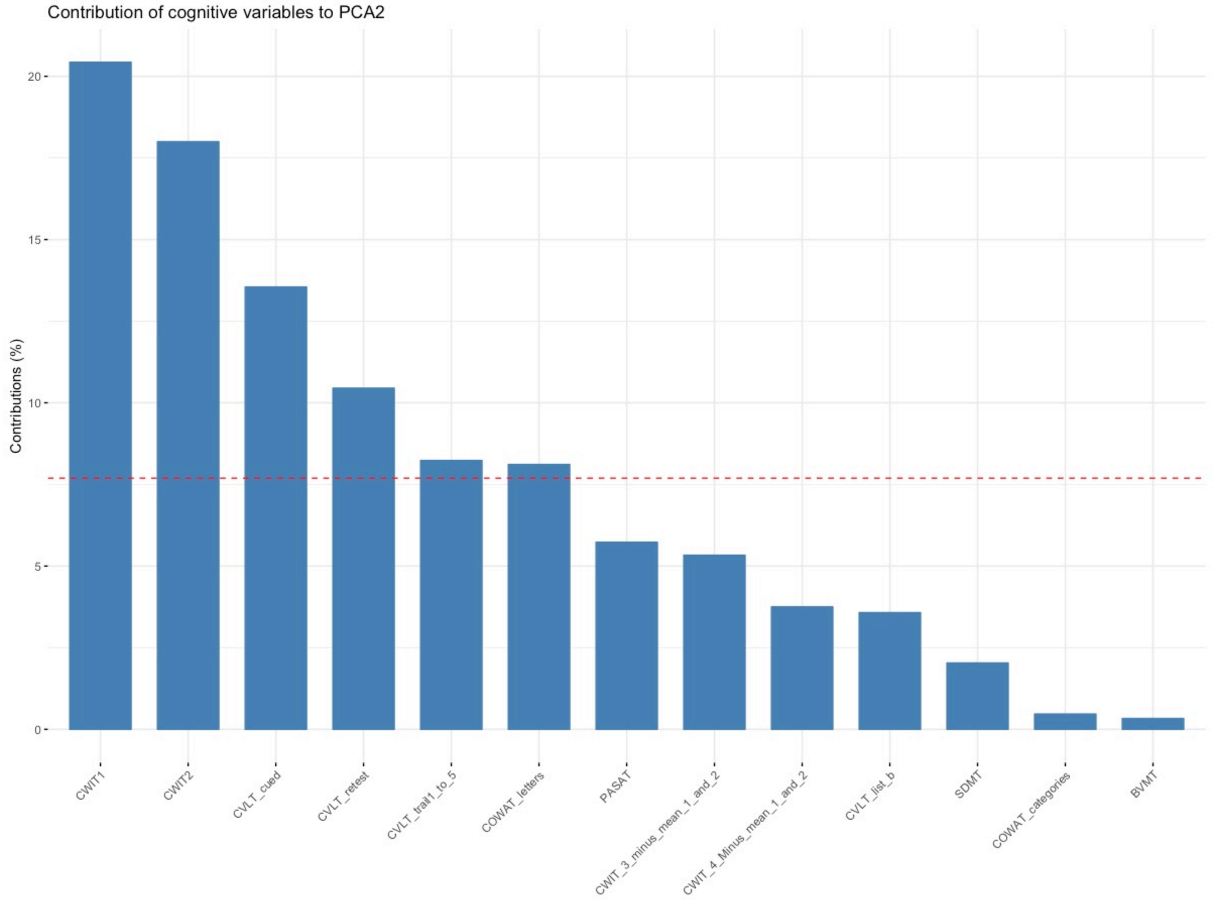


**Supplementary figure 5. Histogram visualizing the individual contributions of the cognitive variables in the second principal component.** CWIT1; D-KEFS CWIT: Color Naming, CWIT2; D-KEFS CWIT: Word Reading, CVLT_cued; CVLT-II: Short-delay cued recall for list A, CVLT_retest; CVLT-II: Short-delay free recall for list A, CVLT_trail_1_to_5; CVLT-II: Immediate free recall for list A, COWAT_letters; COWAT: Letter fluency, PASAT; Paced Auditory Serial Addition Test, CWIT_3_Minus_mean_1_and_2; Inhibition minus mean of Color Naming and Word Reading, CWIT_4_Minus_mean_1_and_2; Inhibition/Switching minus mean of Color Naming and Word Reading, CVLT_list_b; CVLT-II: Immediate free recall for list B, SDMT; Symbol Digit Modalities Test, COWAT_categories; COWAT: Category fluency, BVMT; Brief Visuospatial Memory Test – Revised Edition.

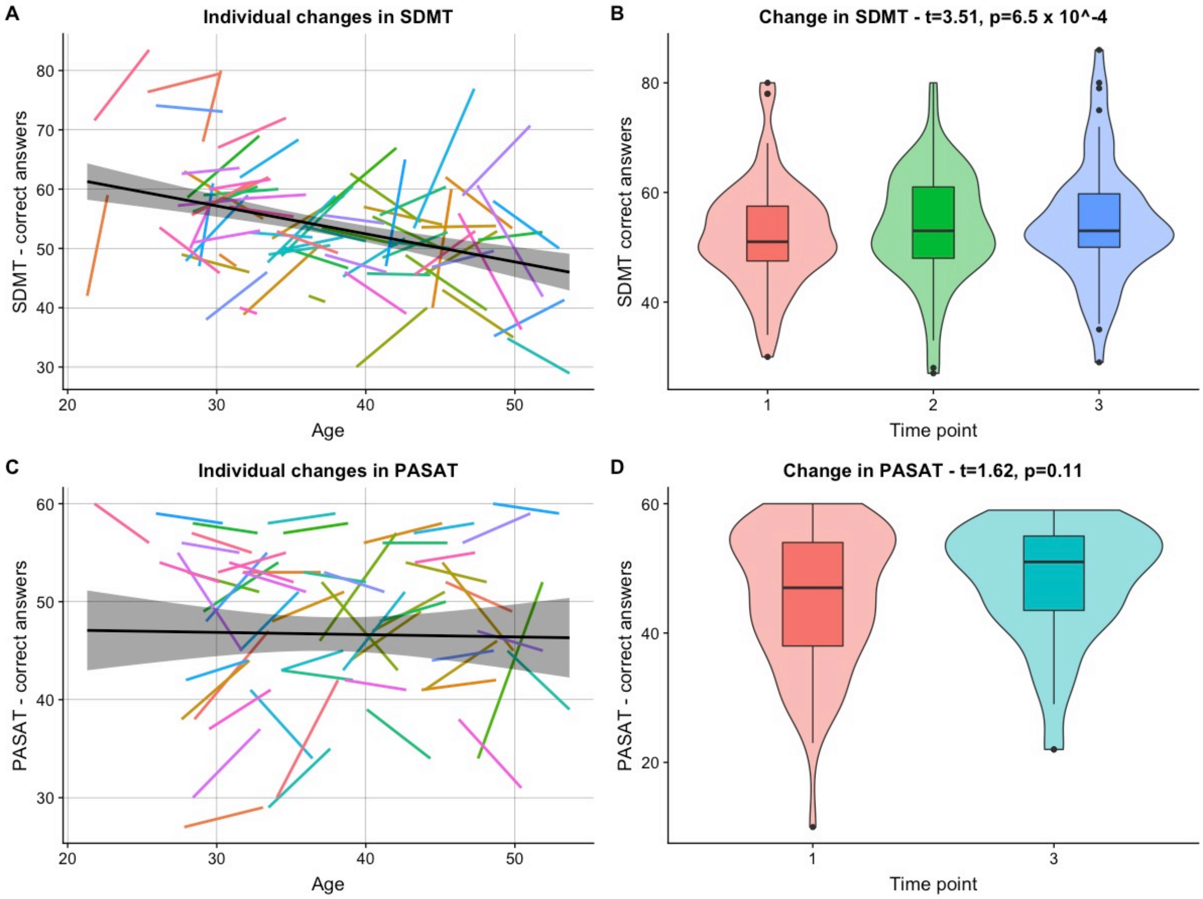


**Supplementary figure 6. Longitudinal results for SDMT and PASAT.** In (A) and (C) the individual regression lines for all subjects are depicted for SDMT and PASAT, respectively. The black lines in (A) and (C) are the summarized regression lines for all data across all time points with the surrounding confidence interval. Furthermore, in (B) and (D) the boxplots for all subjects at all time points are shown for SDMT and PASAT, respectively. SDMT ($t=3.51$, $p=6.5 \times 10^{-4}$) displayed a significant increase in results across all the time points, as measured by a LME model.

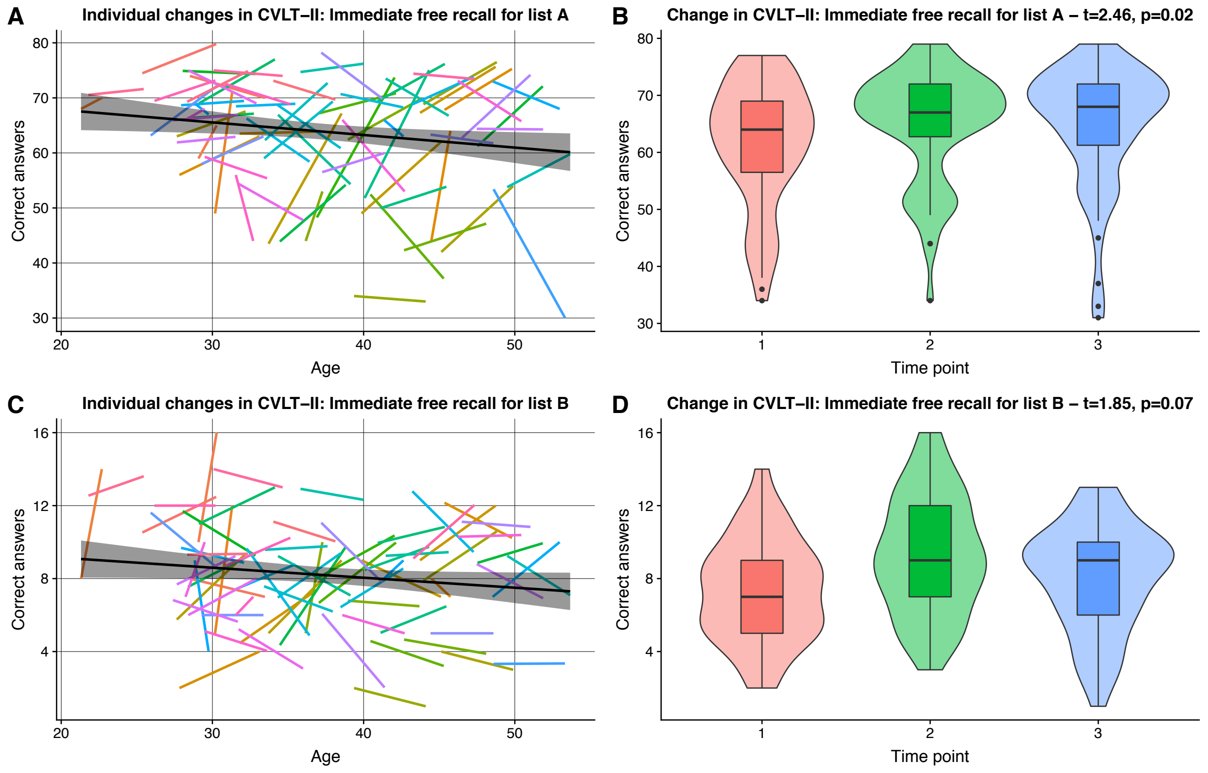


**Supplementary figure 7. Longitudinal results for CVLT-II: Immediate free recall for list A and Immediate free recall for list B.** In (A) and (C) the individual regression lines for all subjects are depicted for CVLT-II: Immediate free recall for list A and Immediate free recall for list B, respectively. The black lines in (A) and (C) are the summarized regression lines for all data across all time points with the surrounding confidence interval. Furthermore, in (B) and (D) the boxplots for all subjects at all time points are shown for CVLT-II: Immediate free recall for list A and Immediate free recall for list B, respectively. CVLT-II: Immediate free recall for list A (t=2.46, p=0.02) displayed a significant increase in results across all the time points, as measured by a LME model.

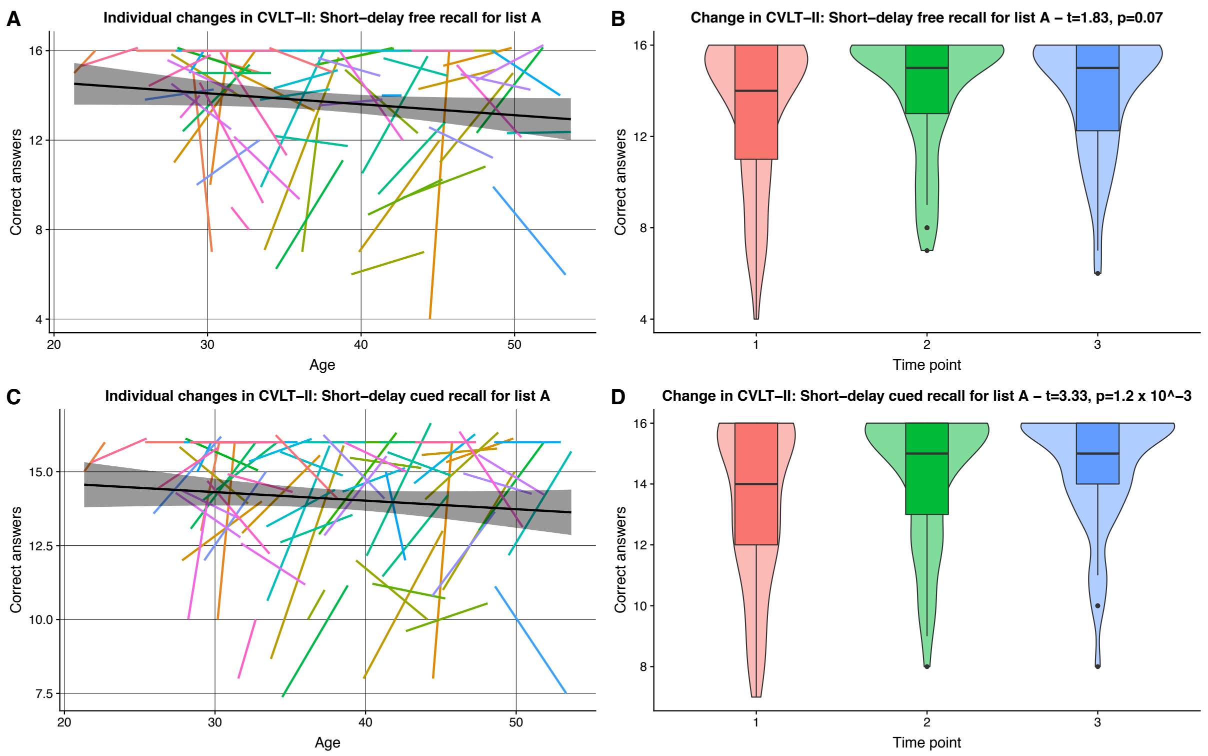


**Supplementary figure 8. Longitudinal results for CVLT-II: Short-delay free recall for list A and Short-delay cued recall for list A.** In (A) and (C) the individual regression lines for all subjects are depicted for both CVLT-II: Short-delay free recall for list A and Short-delay cued recall for list A, respectively. The black lines in (A) and (C) are the summarized regression lines for all data across all time points with the surrounding confidence interval. Furthermore, in (B) and (D) the boxplots for all subjects at all time points are shown for both CVLT-II: Short-delay free recall for list A and Short-delay cued recall for list A, respectively. CVLT-II: Short-delay free recall for list A (t=3.33, p=1.2 x $10^{-3}$) displayed a significant increase in results across all the time points, as measured by a LME model.

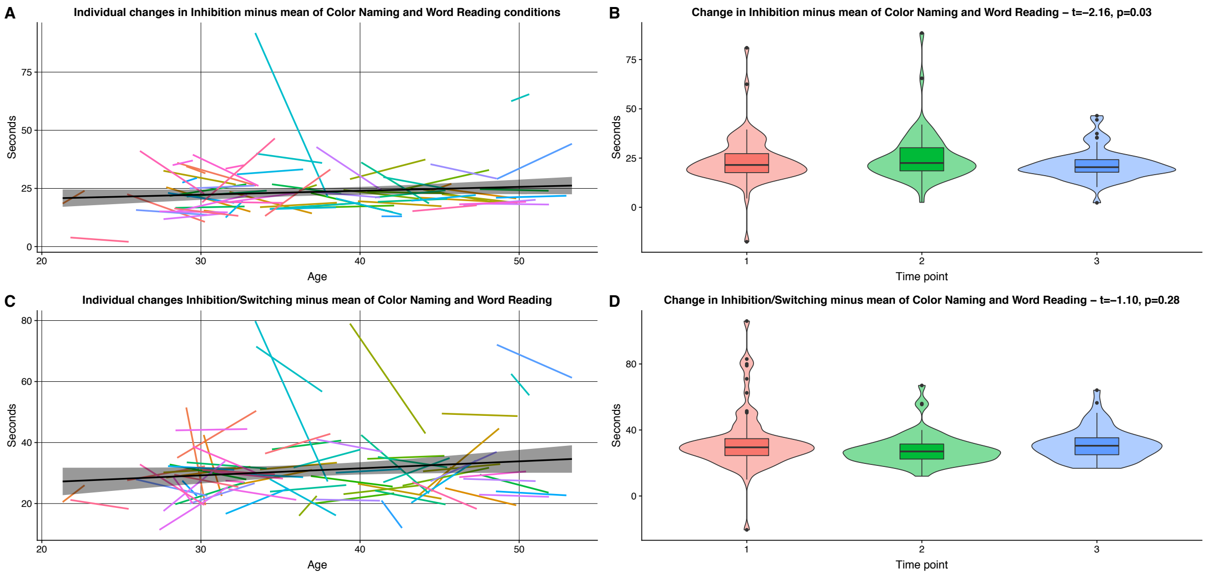


**Supplementary figure 9. Longitudinal results for Inhibition minus mean of Color Naming and Word Reading interactions of CWIT and Inhibition/Switching minus mean of Color Naming and Word Reading conditions of CWIT.** In (A) and (C) the individual regression lines for all subjects are depicted for both Inhibition minus mean of Color Naming and Word Reading interactions of CWIT and Inhibition/Switching minus mean of Color Naming and Word Reading conditions of CWIT, respectively. The black lines in (A) and (C) are the summarized regression lines for all data across all time points with the surrounding confidence interval. Furthermore, in (B) and (D) the boxplots for all subjects at all time points are shown for both Inhibition minus mean of Color Naming and Word Reading interactions of CWIT and Inhibition/Switching minus mean of Color Naming and Word Reading conditions of CWIT, respectively. Inhibition minus mean of Color Naming and Word Reading interactions of CWIT (t=-2.16, p=0.03) displayed a significant decrease in results across all the time points, as measured by a LME model.

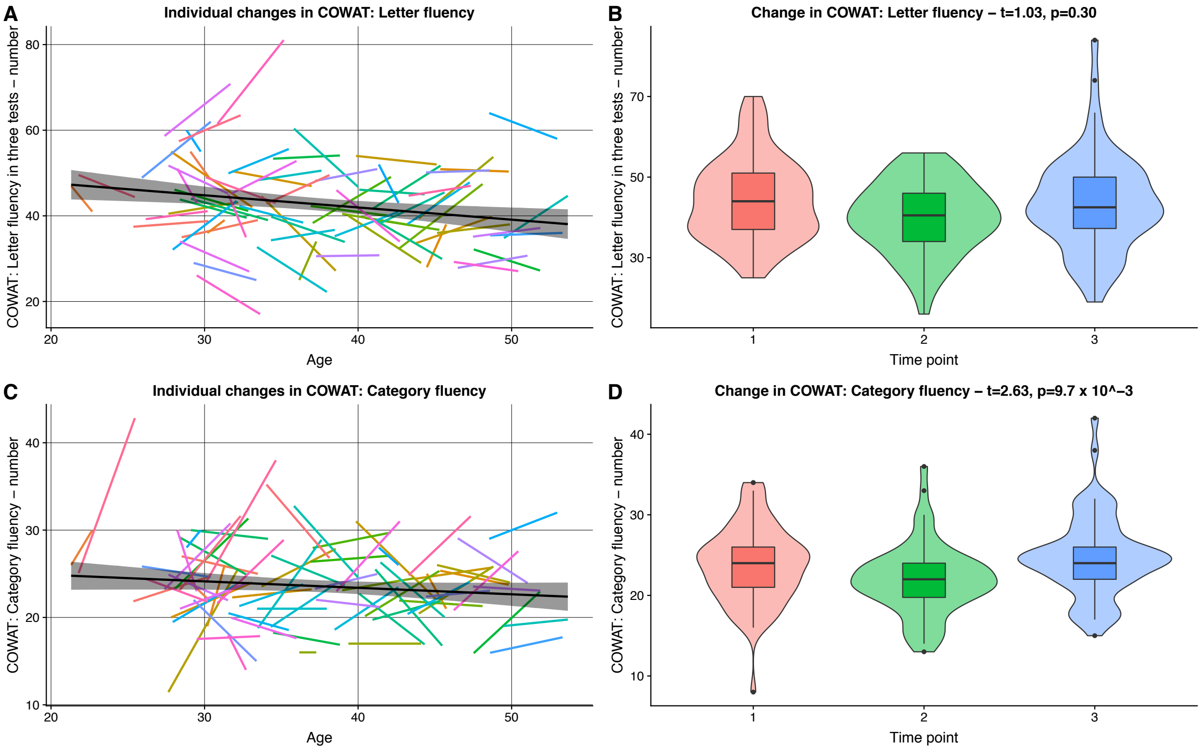


**Supplementary figure 10. Longitudinal results for COWAT: Letter fluency and Category fluency.** In (A) and (C) the individual regression lines for all subjects are depicted for both the three tests of COWAT: Letter fluency and Category fluency, respectively. The black lines in (A) and (C) are the summarized regression lines for all data across all time points with the surrounding confidence interval. Furthermore, in (B) and (D) the boxplots for all subjects at all time points are shown for both the COWAT: Letter fluency and Category fluency three tests, respectively. COWAT: Category fluency ($t=2.63$, $p=9.7 \times 10^{-3}$) displayed a significant increase in results across all the time points, as measured by a LME model.

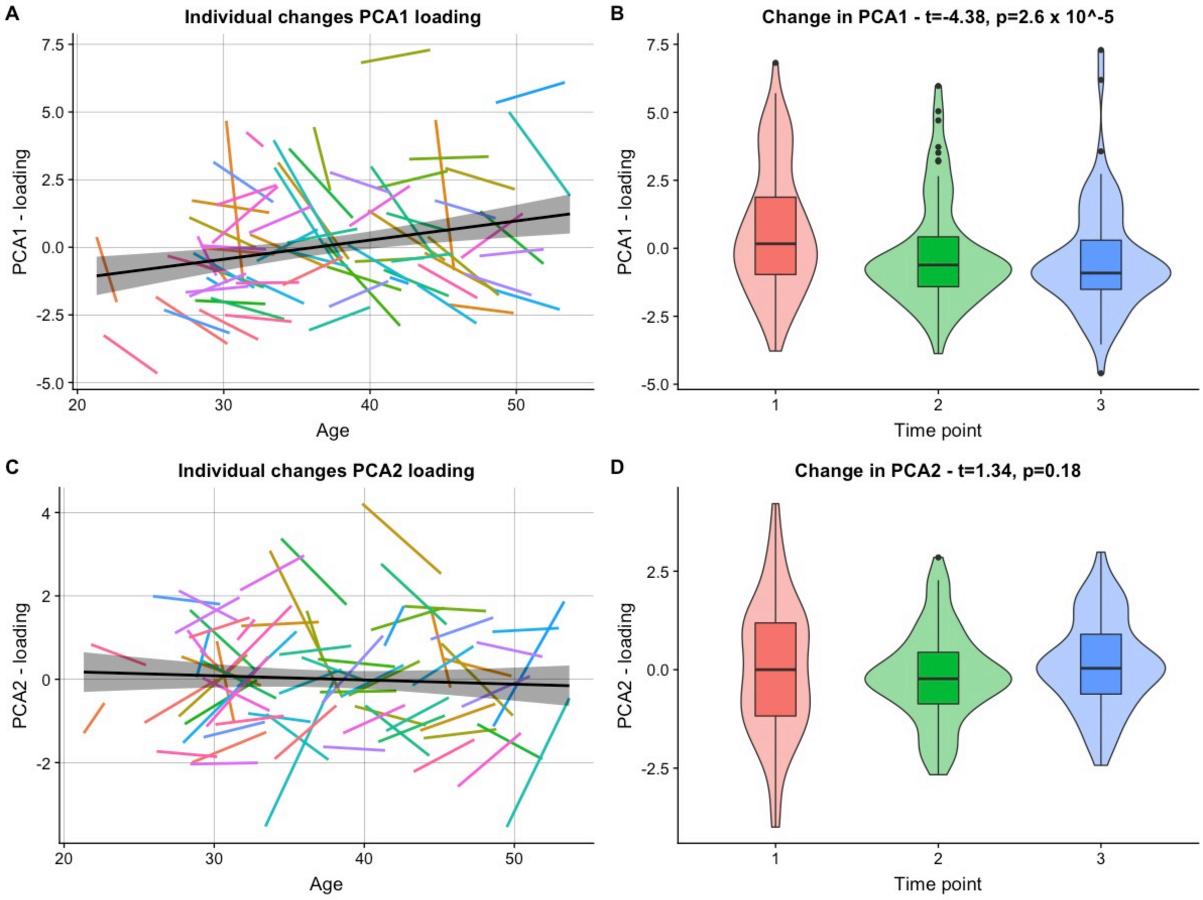


**Supplementary figure 11. Principal component analysis (PCA) - Longitudinal results for loadings of PCA1 and PCA2.** In (A) and (C) the individual regression lines for all subjects are depicted for PCA1 and PCA 2 loadings, respectively. The black lines in (A) and (C) are the summarized regression lines for all data across all time points with the surrounding confidence interval. Furthermore, in (B) and (D) the boxplots for all subjects at all time points are shown for both PCA1 and PCA2 loadings, respectively. PCA1 ($t=-4.38$, $p=2.6 \times 10^{-5}$) displayed a significant decrease in results across all the time points, as measured by a LME model.

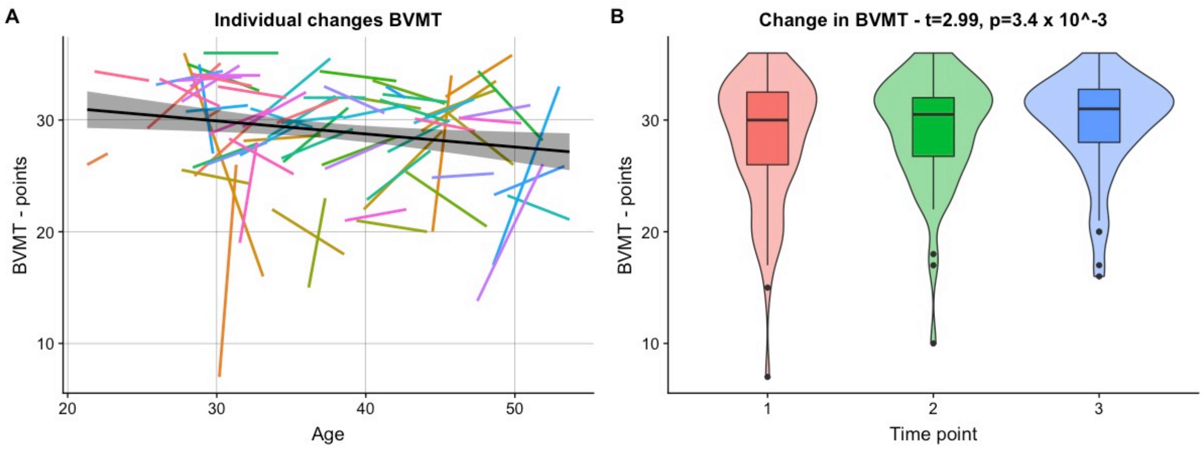


**Supplementary figure 12. Longitudinal results for BVMT.** In (A) the individual regression lines for all subjects are depicted for BVMT. The black line in (A) is the summarized regression line for all data across all time points with the surrounding confidence interval. Furthermore, in (B) the boxplots for all subjects at all time points are shown for BVMT. BVMT (t=2.99, p=3.4 x $10^{-3}$) displayed a significant increase in results across all the time points, as measured by a LME model.

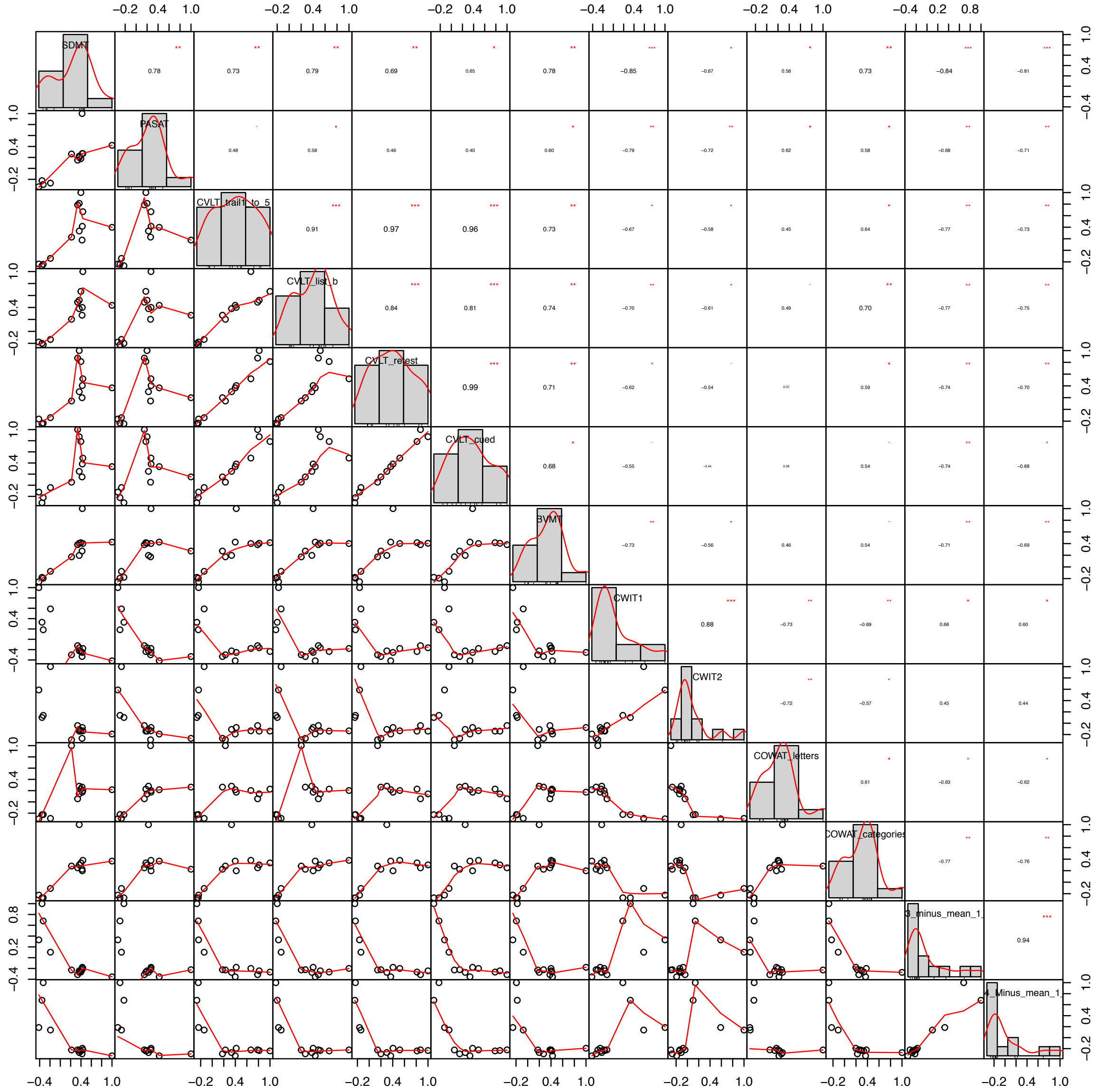


**Supplementary figure 13. Correlation matrix for the cognitive tests.** SDMT; Symbol Digit Modalities Test, PASAT; Paced Auditory Serial Addition Test, CVLT_trail_1_to_5; CVLT-II: Immediate free recall for list A, CVLT_list_b; CVLT-II: Immediate free recall for list B, CVLT_retest; CVLT-II: Short-delay free recall for list A, CVLT_cued; CVLT-II: Short-delay cued recall for list A, BVMT; Brief Visuospatial Memory Test – Revised Edition, CWIT1; D-KEFS CWIT: Color Naming, CWIT2; D-KEFS CWIT: Word Reading, COWAT_letters; COWAT: Letter fluency, COWAT_categories; COWAT: Category fluency, CWIT_3_Minus_mean_1_and_2; Inhibition minus mean of Color Naming and Word Reading, CWIT_4_Minus_mean_1_and_2; Inhibition/Switching minus mean of Color Naming and Word Reading.

### REFERENCES SUPPLEMENTARY MATERIAL


1. Smith A. *Symbol digit modalities test: Manual.* Los Angeles, CA: Western Psychological Services, 1982.
2. Rao SM, Leo GJ, Bernardin L, et al. Cognitive dysfunction in multiple sclerosis. I. Frequency, patterns, and prediction. *Neurology* 1991; 41: 685-691. 1991/05/01.
3. Delis DC, Kaplan E and Kramer JH. *Delis–Kaplan executive function system: Examiner's manual*. San Antonio, TX: The Psychological Corporation, 2001.
4. Delis DC, Kramer JH, Kaplan E, et al. *California verbal learning test – second edition. Adult version. Manual*. San Antonio, TX: Psychological Corporation, 2000.
5. Delis D, Kramer J, Kaplan E, et al. *California Verbal Learning Test (CVLT–II). Norwegian Manual Supplement*. Stockholm: Pearson Assessment, 2004.
6. Benedict RHB. *Brief visuospatial memory test - revised: Professional manual*. Lutz, FL: Psychological Assessment Resources, 1997.
7. Spreen O and Strauss E. *A compendium of neuropsychological tests : administration, norms, and commentary*. 2nd ed. ed. New York: Oxford University Press, 1998.
8. Kaufmann T, van der Meer D, Doan NT, et al. Genetics of brain age suggest an overlap with common brain disorders. *bioRxiv* 2018. DOI: 10.1101/303164.
9. Cole JH and Franke K. Predicting Age Using Neuroimaging: Innovative Brain Ageing Biomarkers. *Trends Neurosci* 2017; 40: 681-690. DOI: 10.1016/j.tins.2017.10.001.
