## Supplementary Tables for "Brain age estimation is a sensitive marker of processing speed in the early course of multiple sclerosis"

**Supplementary table 1.** An overview the resulting components from the principal component analysis run with all the cognitive data as input.

| *PCA component* | *Eigenvalue* | *Total variance explained (%)* | *Cumulative precentage explained (%)* |
|---|---|---|---|
| PCA1 | 4.60 | 35.39 | 35.39 |
| PCA2 | 1.83 | 14.11 | 49.50 |
| PCA3 | 1.19 | 9.13 | 58.62 |
| PCA4 | 0.99 | 7.59 | 66.22 |
| PCA5 | 0.94 | 7.28 | 73.49 |
| PCA6 | 0.75 | 5.79 | 79.28 |
| PCA7 | 0.65 | 5.01 | 84.29 |
| PCA8 | 0.56 | 4.29 | 88.58 |
| PCA9 | 0.49 | 3.74 | 92.32 |
| PCA10 | 0.40 | 3.08 | 95.40 |
| PCA11 | 0.32 | 2.43 | 97.82 |
| PCA12 | 0.15 | 1.17 | 99.00 |
| PCA13 | 0.13 | 1.00 | 100.00 |

PCA - Principal Component Analysis

**Supplementary table 2.** An overview of the intraclass correlation coefficients (ICC) for all the cognitive tests for all time points.

| | ***Time point 1 vs. time point 2*** | | | | ***Time point 2 vs. time point 3*** | | | | ***Time point 1 vs. time point 3*** | | | | ***All time points*** | | | |
|---|---|---|---|---|---|---|---|---|---|---|---|---|---|---|---|---|
| *Neurocognitive tests* | *ICC (95% CI)* | *f-value* | *p* | *n* | *ICC (95% CI)* | *f-value* | *p* | *n* | *ICC (95% CI)* | *f-value* | *p* | *n* | *ICC (95% CI)* | *f-value* | *p* | *n* |
| SDMT | 0.73 (0.59-0.83) | 6.4 | $2.2 \times 10^{-12}$ | 64 | 0.68 (0.51-0.80) | 5.3 | $3.1 \times 10^{-9}$ | 55 | 0.72 (0.57-0.82) | 8.7 | $1.6 \times 10^{-11}$ | 61 | 0.72 (0.61-0.81) | 8.7 | $8.7 \times 10^{-22}$ | 55 |
| PASAT | NA | NA | NA | NA | NA | NA | NA | NA | 0.76 (0.62-0.85) | 7.3 | $2.9 \times 10^{-12}$ | 56 | NA | NA | NA | NA |
| CVLT-II: Immediate free recall for list A | 0.58 (0.39-0.72) | 3.8 | $1.7 \times 10^{-7}$ | 64 | 0.79 (0.66-0.87) | 8.4 | $2.1 \times 10^{-13}$ | 55 | 0.57 (0.37-0.72) | 3.6 | $6.0 \times 10^{-7}$ | 61 | 0.65 (0.51-0.76) | 6.5 | $6.4 \times 10^{-17}$ | 55 |
| CVLT-II: Immediate free recall for list B | 0.31 (0.08-0.52) | 1.9 | $5.2 \times 10^{-3}$ | 64 | 0.63 (0.44-0.76) | 4.4 | $9.9 \times 10^{-8}$ | 55 | 0.64 (0.47-0.77) | 4.6 | $6.4 \times 10^{-9}$ | 61 | 0.54 (0.39-0.68) | 4.5 | $1.2 \times 10^{-11}$ | 55 |
| CVLT-II: Short-delay free recall for list A | 0.41 (0.19-0.60) | 2.4 | $3.0 \times 10^{-4}$ | 64 | 0.52 (0.29-0.69) | 3.1 | $2.1 \times 10^{-5}$ | 55 | 0.54 (0.34-0.70) | 3.4 | $2.6 \times 10^{-6}$ | 61 | 0.54 (0.39-0.68) | 4.5 | $1.2 \times 10^{-11}$ | 55 |
| CVLT-II: Short-delay cued recall for list A | 0.37 (0.14-0.56) | 2.2 | $1.2 \times 10^{-3}$ | 64 | 0.57 (0.36-0.72) | 3.7 | $1.9 \times 10^{-6}$ | 55 | 0.36 (0.12-0.56) | 2.1 | $2.0 \times 10^{-3}$ | 61 | 0.43 (0.27-0.59) | 3.3 | $6.0 \times 10^{-8}$ | 55 |
| BVMT-R | 0.50 (0.30-0.67) | 3.0 | $8.7 \times 10^{-6}$ | 64 | 0.61 (0.41-0.75) | 4.1 | $3.0 \times 10^{-7}$ | 55 | 0.50 (0.29-0.67) | 3.0 | $1.6 \times 10^{-5}$ | 61 | 0.64 (0.50-0.75) | 6.3 | $2.3 \times 10^{-16}$ | 55 |
| D-KEFS CWIT: Color Naming | 0.73 (0.60-0.83) | 6.5 | $1.2 \times 10^{-12}$ | 64 | 0.64 (0.45-0.77) | 4.5 | $7.1 \times 10^{-8}$ | 54 | 0.66 (0.49-0.78) | 4.8 | $3.6 \times 10^{-9}$ | 60 | 0.66 (0.53-0.77) | 7.0 | $1.1 \times 10^{-17}$ | 54 |
| D-KEFS CWIT: Word Reading | 0.71 (0.56-0.81 | 5.9 | $1.5 \times 10^{-11}$ | 64 | 0.72 (0.56-0.82) | 6.1 | $1.9 \times 10^{-10}$ | 55 | 0.70 (0.54-0.81) | 5.6 | $1.0 \times 10^{-10}$ | 61 | 0.72 (0.61-0.82) | 8.9 | $3.9 \times 10^{-22}$ | 55 |
| D-KEFS CWIT: Inhibition | 0.56 (0.37-0.71) | 3.6 | $5.9 \times 10^{-7}$ | 63 | 0.85 (0.75-0.91) | 12.2 | $1.6 \times 10^{-16}$ | 53 | 0.42 (0.18-0.61) | 2.4 | $4.8 \times 10^{-4}$ | 58 | 0.55 (0.39-0.69) | 4.7 | $1.4 \times 10^{-11}$ | 52 |
| D-KEFS CWIT: Inhibition/Switching | 0.55 (0.35-0.70) | 3.4 | $1.2 \times 10^{-6}$ | 63 | 0.78 (0.65-0.87) | 8.2 | $5.9 \times 10^{-13}$ | 54 | 0.59 (0.40-0.73) | 3.9 | $2.8 \times 10^{-7}$ | 59 | 0.61 (0.46-0.73) | 5.7 | $2.4 \times 10^{-14}$ | 53 |
| *Inhibition minus mean of Color Naming and Word Reading* | 0.97 (0.96-0.98) | 76.8 | $9.9 \times 10^{-43}$ | 63 | 0.27 (-0.00-0.50 | 1.7 | 0.03 | 52 | 0.28 (0.03-0.50) | 1.8 | 0.01 | 58 | 0.57 (0.43-0.71) | 5.1 | $1.0 \times 10^{-12}$ | 52 |
| *Inhibition/Switching minus mean of Color Naming and Word Reading* | 0.45 (0.23-0.63) | 2.6 | $8.7 \times 10^{-5}$ | 63 | 0.65 (0.46-0.78) | 4.7 | $3.7 \times 10^{-8}$ | 54 | 0.54 (0.33-0.70) | 3.3 | $4.0 \times 10^{-6}$ | 59 | 0.53 (0.38-0.68) | 4.4 | $4.6 \times 10^{-11}$ | 53 |
| COWAT: Letter fluency | 0.56 (0.37-0.71) | 3.5 | $5.7 \times 10^{-7}$ | 64 | 0.59 (0.38-0.74) | 3.8 | $9.1 \times 10^{-7}$ | 55 | 0.76 (0.64-0.85) | 7.5 | $1.6 \times 10^{-13}$ | 61 | 0.64 (0.50-0.75) | 6.3 | $2.6 \times 10^{-16}$ | 55 |
| COWAT: Category fluency | 0.37 (0.14-0.56) | 2.2 | $1.1 \times 10^{-3}$ | 64 | 0.34 (0.09-0.55) | 2.0 | $4.7 \times 10^{-3}$ | 55 | 0.27 (0.03-0.49) | 0.3 | 0.015 | 61 | 0.32 (0.15-0.49) | 2.4 | $5.4 \times 10^{-5}$ | 55 |

ICC - Intraclass Correlation Coefficient, CI - Confidence Interval, SDMT - Symbol Digital Modalities Test, PASAT - Paced Auditory Serial Addition Test, CVLT-II - California Verbal Learning Test-II, BVMT-R - Brief Visuospatial Memory Test Revised , CWIT - Color Word Interference Test , COWAT - Controlled Oral Word Association test

**Supplementary table 3.** An overview of the intraclass correlation coefficients (ICC) for all the components from the principal component analysis (PCA) at all time points.

| | ***Time point 1 vs. time point 2*** | | | | ***Time point 2 vs. time point 3*** | | | | ***Time point 1 vs. time point 3*** | | | | ***All time points*** | | | |
|---|---|---|---|---|---|---|---|---|---|---|---|---|---|---|---|---|
| *Principal Component Analysis* | *ICC (95% CI)* | *f-value* | *p* | *n* | *ICC (95% CI)* | *f-value* | *p* | *n* | *ICC (95% CI)* | *f-value* | *p* | *n* | *ICC (95% CI)* | *f-value* | *p* | *n* |
| PC1 | 0.64 (0.48-0.75) | 4.5 | $1.8 \times 10^{-10}$ | 76 | 0.78 (0.68-0.86) | 8.2 | $1.5 \times 10^{-17}$ | 76 | 0.55 (0.37-0.69) | 3.4 | $1.1 \times 10^{-7}$ | 76 | 0.65 (0.54-0.75) | 6.6 | $6.6 \times 10^{-23}$ | 76 |
| PC2 | 0.64 (0.49-0.76) | 4.6 | 1.6 x 10-10 | 76 | 0.60 (0.44-0.73) | 4.0 | $3.1 \times 10^{-9}$ | 76 | 0.61 (0.45-0.74) | 4.2 | $1.3 \times 10^{-9}$ | 76 | 0.62 (0.50-0.72) | 5.9 | $1.1 \times 10^{-20}$ | 76 |
| PC3 | 0.26 (0.04-0.46) | 1.7 | 0.01 | 76 | 0.37 (0.15-0.54 | 2.2 | $5.3 \times 10^{-4}$ | 76 | 0.11 (-0.11-0.33) | 1.3 | 0.17 | 76 | 0.24 (0.10-0.39) | 1.9 | $3.6 \times 10^{-4}$ | 76 |
| PC4 | 0.20 (-0.02-0.41) | 1.5 | 0.04 | 76 | 0.20 (-0.02-0.41) | 1.5 | 0.04 | 76 | 0.23 (0.00-0.43) | 1.6 | 0.02 | 76 | 0.23 (0.10-0.39) | 1.9 | $3.5 \times 10^{-4}$ | 76 |
| PC5 | 0.32 (0.11-0.51) | 1.9 | $2.2 \times 10^{-3}$ | 76 | 0.24 (0.02-0.44) | 1.6 | 0.02 | 76 | 0.27 (0.05-0.47) | 1.7 | $8.6 \times 10^{-3}$ | 76 | 0.28 (0.14-0.43) | 2.2 | $2.2 \times 10^{-5}$ | 76 |
| PC6 | 0.35 (0.13-0.53) | 2.1 | $1.0 \times 10^{-3}$ | 76 | 0.14 (-0.09-0.35) | 1.3 | 0.11 | 76 | 0.41 (0.21-0.58) | 2.4 | $8.7 \times 10^{-5}$ | 76 | 0.32 (0.18-0.47) | 2.4 | $2.5 \times 10^{-6}$ | 76 |
| PC7 | 0.30 (0.08-0.49) | 1.9 | $3.9 \times 10^{-3}$ | 76 | 0.29 (0.07-0.48) | 1.8 | $5.5 \times 10^{-3}$ | 76 | 0.32 (0.11-0.51) | 2.0 | $2.0 \times 10^{-3}$ | 76 | 0.31 (0.16-0.45) | 2.3 | $5.9 \times 10^{-6}$ | 76 |
| PC8 | -0.08 (-0.30-0.15) | 0.9 | 0.76 | 76 | 0.02 (-0.21-0.24) | 1.0 | 0.44 | 76 | 0.34 (0.13-0.52) | 2.0 | $1.2 \times 10^{-3}$ | 76 | 0.10 (-0.03-0.25) | 1.3 | 0.07 | 76 |
| PC9 | 0.32 (0.10-0.50 | 1.9 | $2.5 \times 10^{-3}$ | 76 | 0.18 (-0.05-0.39) | 1.4 | 0.06 | 76 | 0.28 (0.06-0.47) | 1.8 | $7.1 \times 10^{-3}$ | 76 | 0.26 (0.12-0.41) | 2.1 | $7.4 \times 10^{-5}$ | 76 |
| PC10 | 0.34 (0.13-0.53) | 2.0 | $1.1 \times 10^{-3}$ | 76 | 0.06 (-0.17-0.28) | 1.1 | 0.31 | 76 | 0.20 (-0.03-0.41) | 1.5 | 0.04 | 76 | 0.21 (0.07-0.36) | 1.8 | $1.1 \times 10^{-3}$ | 76 |
| PC11 | 0.49 (0.30-0.65) | 3.0 | $2.2 \times 10^{-6}$ | 76 | 0.11 (-0.12-0.33) | 1.3 | 0.17 | 76 | 0.07 (-0.16-0.29) | 1.2 | 0.28 | 76 | 0.24 (0.10-0.39) | 2.0 | $2.5 \times 10^{-4}$ | 76 |
| PC12 | 0.03 (-0.20-0.25) | 1.1 | 0.40 | 76 | -0.06 (-0.28-0.16) | 0.9 | 0.71 | 76 | 0.18 (-0.04-0.39) | 1.4 | 0.06 | 76 | 0.07 (-0.06-0.21) | 1.2 | 0.16 | 76 |
| PC13 | -0.30 (-0.49- -0.08) | 0.5 | 1.00 | 76 | 0.06 (-0.16-0.28) | 1.1 | 0.30 | 76 | -0.03 (-0.25-0.19) | 0.9 | 0.61 | 76 | -0.09 (-0.19-0.04) | 0.8 | 0.92 | 76 |

ICC - Intraclass Correlation Coefficient, CI - Confidence Interval, PCA - Principal Component Analysis, PC - Principal Component

**Supplementary table 4.** An overview of all correlations between the cognitive tests and MRI variables in the cohort. All longitudinal tests were run in R using the "nlme" package to calculate LME models, also accounting for months since time point 1, age and gender as fixed parameters. The linear models are calculated using the "stats" package in R, also accounting age and gender as fixed parameters. Brain age gaps are corrected for age and gender effects. Significant associations are marked as bold, while the associations still significant after correcting for multiple comparisons are additionally marked as red.

| | **MRI variables** | | | | | | | | | | | | | | | | | |
|---|---|---|---|---|---|---|---|---|---|---|---|---|---|---|---|---|---|---|
| | *Brain age gap* | | *Brain volume* | | *White matter volume* | | *Grey matter volume* | | *Thalamus volume* | | *Normalized brain volume* | | *Normalized white matter volume* | | *Normalized grey matter volume* | | *Normalized thalamus volume* | |
| **Test variables** | t | p | t | p | t | p | t | p | t | p | t | p | t | p | t | p | t | p |
| *SDMT* | | | | | | | | | | | | | | | | | | |
| Time point 1 | -1.58 | 0.12 | 1.53 | 0.13 | 1.02 | 0.31 | 1.91 | 0.06 | 1.49 | 0.14 | -1.21 | 0.23 | -1.29 | 0.20 | -0.73 | 0.47 | 0.09 | 0.93 |
| Time point 2 | -0.48 | 0.63 | 0.66 | 0.51 | 0.34 | 0.74 | 0.91 | 0.37 | 0.67 | 0.51 | 0.19 | 0.85 | -0.06 | 0.95 | 0.32 | 0.75 | 0.39 | 0.70 |
| Time point 3 | -0.39 | 0.70 | 1.35 | 0.18 | 1.69 | 0.10 | 0.84 | 0.41 | 1.85 | 0.07 | 0.54 | 0.59 | 1.58 | 0.12 | -0.76 | 0.45 | 1.4 | 0.16 |
| All time points | -0.37 | 0.71 | 1.35 | 0.18 | 1.46 | 0.15 | 1.02 | 0.31 | 1.73 | 0.09 | -0.40 | 0.69 | 0.42 | 0.67 | -1.00 | 0.32 | 0.85 | 0.39 |
| *PASAT* | | | | | | | | | | | | | | | | | | |
| Time point 1 | -0.74 | 0.46 | -1.03 | 0.30 | -1.70 | 0.09 | -0.28 | 0.78 | 0.04 | 0.97 | 0.55 | 0.59 | -1.06 | 0.29 | 1.93 | 0.06 | 1.04 | 0.30 |
| Time point 3 | 0.33 | 0.75 | 0.87 | 0.39 | 1.17 | 0.25 | 0.42 | 0.68 | 0.66 | 0.51 | 0.55 | 0.59 | 1.27 | 0.21 | -0.49 | 0.62 | 0.40 | 0.69 |
| All time points | -0.71 | 0.48 | 0.22 | 0.83 | -0.21 | 0.84 | 0.55 | 0.58 | 0.70 | 0.49 | 0.22 | 0.83 | -0.50 | 0.62 | 0.66 | 0.51 | 0.82 | 0.42 |
| *CVLT-II: Immediate free recall for list A* | | | | | | | | | | | | | | | | | | |
| Time point 1 | 0.34 | 0.73 | 1.29 | 0.20 | 1.51 | 0.14 | 0.98 | 0.33 | 0.99 | 0.33 | -1.06 | 0.29 | 0.07 | 0.94 | -1.71 | 0.09 | -0.38 | 0.71 |
| Time point 2 | 1.11 | 0.27 | -0.88 | 0.38 | -0.66 | 0.52 | -1.00 | 0.32 | 0.49 | 0.63 | -0.94 | 0.35 | -0.59 | 0.56 | -0.90 | 0.37 | 0.83 | 0.41 |
| Time point 3 | -0.27 | 0.79 | 1.63 | 0.11 | **2.24** | **0.03** | 0.83 | 0.41 | **3.1** | **2.8 x $10^{-3}$** | -0.49 | 0.63 | 1.30 | 0.20 | **-2.15** | **0.04** | **2.08** | **0.04** |
| All time points | 0.23 | 0.82 | 1.19 | 0.23 | 1.65 | 0.10 | 0.62 | 0.53 | **1.94** | **0.05** | -0.93 | 0.35 | 0.48 | 0.64 | -1.85 | 0.07 | 0.91 | 0.36 |
| *CVLT-II: Immediate free recall for list B* | | | | | | | | | | | | | | | | | | |
| Time point 1 | -1.12 | 0.27 | **2.23** | **0.03** | **2.32** | **0.02** | **1.99** | **0.05** | **3.04** | **3.4 x $10^{-3}$** | -0.05 | 0.96 | 0.94 | 0.35 | -0.90 | 0.37 | 1.80 | 0.08 |
| Time point 2 | 1.05 | 0.30 | 0.08 | 0.94 | 0.61 | 0.55 | -0.52 | 0.61 | 0.87 | 0.39 | -1.07 | 0.29 | 0.16 | 0.88 | -1.92 | 0.06 | 0.42 | 0.68 |
| Time point 3 | 0.64 | 0.53 | **2.34** | **0.02** | **2.96** | **4.5 x $10^{-3}$** | 1.46 | 0.15 | **2.69** | **0.01** | -0.60 | 0.55 | 1.39 | 0.17 | **-2.45** | **0.02** | 0.95 | 0.35 |
| All time points | 0.34 | 0.73 | **2.01** | **0.05** | **2.44** | **0.02** | 1.37 | 0.17 | **2.71** | **7.8 x $10^{-3}$** | -0.72 | 0.47 | 0.87 | 0.38 | **-1.99** | **0.05** | 1.24 | 0.22 |
| *CVLT-II: Short-delay free recall for list A* | | | | | | | | | | | | | | | | | | |
| Time point 1 | 0.58 | 0.56 | 0.04 | 0.97 | 0.14 | 0.89 | 0.00 | 1.00 | 0.77 | 0.45 | -1.43 | 0.16 | -0.79 | 0.44 | -1.38 | 0.17 | 0.28 | 0.78 |
| Time point 2 | 1.79 | 0.08 | **-1.98** | **0.05** | -1.22 | 0.23 | **-2.55** | **0.01** | -0.80 | 0.43 | -0.85 | 0.40 | -0.05 | 0.96 | -1.23 | 0.22 | 0.16 | 0.88 |
| Time point 3 | -0.10 | 0.92 | 0.88 | 0.38 | 1.51 | 0.14 | 0.15 | 0.88 | **2.98** | **4.3 x $10^{-3}$** | 0.38 | 0.71 | 1.75 | 0.09 | -1.11 | 0.27 | **3.13** | **2.8 x $10^{-3}$** |
| All time points | 1.11 | 0.27 | -0.16 | 0.87 | 0.30 | 0.77 | -0.55 | 0.58 | 0.99 | 0.32 | -1.10 | 0.27 | -0.00 | 1.00 | -1.61 | 0.11 | 0.91 | 0.36 |
| *CVLT-II: Short-delay cued recall for list A* | | | | | | | | | | | | | | | | | | |
| Time point 1 | 0.64 | 0.53 | -0.66 | 0.51 | -0.39 | 0.70 | -0.83 | 0.41 | -0.28 | 0.78 | -0.66 | 0.51 | -0.13 | 0.89 | -0.92 | 0.42 | -0.12 | 0.90 |
| Time point 2 | 1.17 | 0.25 | -0.86 | 0.40 | -0.57 | 0.57 | -1.02 | 0.31 | 0.86 | 0.39 | -0.81 | 0.42 | -0.36 | 0.72 | -0.89 | 0.38 | 1.26 | 0.21 |
| Time point 3 | -0.26 | 0.80 | 0.57 | 0.57 | 1.06 | 0.29 | 0.05 | 0.96 | **2.93** | **4.9 x $10^{-3}$** | 0.42 | 0.68 | 1.37 | 0.18 | -0.62 | 0.54 | **3.3** | **1.6 x $10^{-3}$** |
| All time points | 0.99 | 0.33 | -0.48 | 0.64 | -0.02 | 0.98 | -0.83 | 0.41 | 0.97 | 0.34 | -0.63 | 0.53 | 0.22 | 0.83 | -1.06 | 0.29 | 1.30 | 0.20 |
| *BVMT-R* | | | | | | | | | | | | | | | | | | |
| Time point 1 | -1.15 | 0.25 | 0.76 | 0.45 | -0.13 | 0.90 | 0.92 | 0.36 | 1.39 | 0.17 | -0.17 | 0.87 | -0.13 | 0.90 | -0.09 | 0.93 | 0.5 | 0.30 |
| Time point 2 | 0.39 | 0.70 | 1.28 | 0.21 | 0.63 | 0.53 | 1.04 | 0.30 | 1.41 | 0.16 | 0.03 | 0.98 | 0.63 | 0.53 | -0.49 | 0.62 | 0.73 | 0.47 |
| Time point 3 | -0.10 | 0.92 | 1.91 | 0.06 | **2.51** | **0.01** | 1.15 | 0.26 | **2.76** | **7.9 x $10^{-3}$** | -0.31 | 0.76 | 1.45 | 0.15 | -1.90 | 0.06 | 1.69 | 0.10 |
| All time points | -0.70 | 0.49 | 1.75 | 0.08 | 1.81 | 0.07 | 1.50 | 0.14 | 1.84 | 0.07 | 0.11 | 0.91 | 0.71 | 0.48 | -0.43 | 0.67 | 0.81 | 0.42 |
| *D-KEFS CWIT: Color Naming* | | | | | | | | | | | | | | | | | | |
| Time point 1 | 0.56 | 0.58 | -0.40 | 0.69 | -0.19 | 0.85 | -0.60 | 0.55 | -1.26 | 0.21 | -0.86 | 0.39 | -0.37 | 0.71 | -1.03 | 0.31 | -1.79 | 0.32 |
| Time point 2 | 0.82 | 0.42 | 1.00 | 0.32 | 1.30 | 0.20 | 0.58 | 0.56 | 0.07 | 0.95 | -1.02 | 0.31 | 0.04 | 0.97 | -1.69 | 0.10 | -1.31 | 0.20 |
| Time point 3 | 0.73 | 0.47 | -1.73 | 0.09 | -1.79 | 0.08 | -1.46 | 0.15 | **-2.16** | **0.036** | 0.35 | 0.73 | -0.53 | 0.60 | 1.12 | 0.27 | -1.00 | 0.32 |
| All time points | **2.84** | **5.4 x $10^{-3}$** | -1.05 | 0.29 | -0.64 | 0.53 | -1.31 | 0.19 | -1.75 | 0.08 | -1.65 | 0.10 | -0.87 | 0.38 | -1.60 | 0.12 | **-2.20** | **0.03** |
| *D-KEFS CWIT: Word Reading* | | | | | | | | | | | | | | | | | | |
| Time point 1 | 1.36 | 0.18 | **-2.23** | **0.03** | **-2.08** | **0.04** | **-2.28** | **0.03** | **-3.28** | **1.6 x $10^{-3}$** | 1.10 | 0.28 | 0.37 | 0.71 | 1.25 | 0.21 | -1.64 | 0.11 |
| Time point 2 | 1.66 | 0.10 | 0.17 | 0.86 | 0.22 | 0.83 | 0.05 | 0.96 | -0.79 | 0.43 | -0.34 | 0.74 | -0.03 | 0.98 | -0.59 | 0.56 | -1.31 | 0.20 |
| Time point 3 | 0.79 | 0.43 | **-2.24** | **0.03** | **-2.03** | **0.05** | **-2.22** | **0.03** | **-3.02** | **3.8 x $10^{-3}$** | 0.69 | 0.49 | 0.13 | 0.89 | 0.91 | 0.37 | -1.57 | 0.12 |
| All time points | 1.36 | 0.18 | **-1.99** | **0.05** | -1.90 | 0.06 | -1.79 | 0.08 | **-3.09** | **2.5 x $10^{-3}$** | 0.38 | 0.71 | -0.11 | 0.91 | 0.55 | 0.58 | -1.93 | 0.06 |
| *D-KEFS CWIT: Inhibition* | | | | | | | | | | | | | | | | | | |
| Time point 1 | 0.98 | 0.33 | -0.87 | 0.39 | -0.97 | 0.33 | -0.68 | 0.50 | -0.97 | 0.33 | -0.53 | 0.60 | -0.94 | 0.35 | 0.04 | 0.97 | -0.83 | 0.41 |
| Time point 2 | 0.52 | 0.61 | -0.55 | 0.59 | -0.80 | 0.43 | -0.21 | 0.84 | -0.93 | 0.36 | -0.11 | 0.91 | -0.76 | 0.45 | 0.54 | 0.59 | -0.83 | 0.41 |
| Time point 3 | 1.11 | 0.27 | -1.52 | 0.13 | -1.49 | 0.14 | -1.37 | 0.18 | **-2.48** | **0.02** | -0.65 | 0.52 | -0.98 | 0.33 | 0.03 | 0.97 | -1.95 | 0.06 |
| All time points | 1.14 | 0.26 | -0.98 | 0.33 | -1.62 | 0.11 | -0.18 | 0.86 | -1.11 | 0.27 | -0.66 | 0.51 | **-1.96** | **0.05** | 0.78 | 0.45 | -0.92 | 0.36 |
| *D-KEFS CWIT: Inhibition/Switching* | | | | | | | | | | | | | | | | | | |
| Time point 1 | 1.67 | 0.10 | -0.83 | 0.41 | -0.67 | 0.51 | -0.90 | 0.37 | -1.49 | 0.14 | -0.69 | 0.49 | -0.50 | 0.62 | -0.57 | 0.57 | -1.58 | 0.12 |
| Time point 2 | -0.10 | 0.92 | **2.17** | **0.03** | **2.33** | **0.02** | 1.75 | 0.09 | 0.45 | 0.65 | 0.11 | 0.91 | 0.90 | 0.37 | -0.67 | 0.51 | -1.06 | 0.29 |
| Time point 3 | 0.27 | 0.79 | -0.71 | 0.48 | -0.79 | 0.43 | -0.51 | 0.61 | -1.83 | 0.07 | 0.07 | 0.94 | -0.43 | 0.67 | 0.61 | 0.55 | -1.58 | 0.12 |
| All time points | 0.61 | 0.54 | -0.43 | 0.67 | -0.54 | 0.59 | -0.21 | 0.83 | -0.97 | 0.33 | -0.12 | 0.90 | -0.50 | 0.62 | 0.33 | 0.74 | -0.94 | 0.35 |
| *Inhibition minus mean of Color Naming and Word Reading* | | | | | | | | | | | | | | | | | | |
| Time point 1 | 0.71 | 0.48 | -0.44 | 0.66 | 1.69 | 0.09 | -0.17 | 0.87 | -0.20 | 0.84 | 0.61 | 0.55 | -1.03 | 0.31 | 0.05 | 0.96 | -0.22 | 0.83 |
| Time point 2 | 0.22 | 0.83 | -0.83 | 0.41 | -1.18 | 0.24 | -0.35 | 0.73 | -0.92 | 0.36 | 0.10 | 0.92 | -0.85 | 0.40 | 0.98 | 0.33 | -0.48 | 0.63 |
| Time point 3 | 1.07 | 0.29 | -0.89 | 0.38 | -0.86 | 0.39 | -0.81 | 0.42 | -1.66 | 0.10 | -1.09 | 0.28 | -1.07 | 0.29 | -0.54 | 0.59 | -1.67 | 0.10 |
| All time points | 0.60 | 0.55 | -0.53 | 0.60 | -1.14 | 0.26 | 0.21 | 0.83 | -0.32 | 0.75 | -0.61 | 0.54 | -1.78 | 0.08 | 0.75 | 0.46 | -0.28 | 0.78 |
| *Inhibition/Switching minus mean of Color Naming and Word Reading* | | | | | | | | | | | | | | | | | | |
| Time point 1 | 1.50 | 0.14 | -0.52 | 0.61 | -0.40 | 0.19 | -0.56 | 0.58 | -0.97 | 0.33 | -0.74 | 0.46 | -0.52 | 0.61 | -0.61 | 0.54 | -1.20 | 0.24 |
| Time point 2 | -0.63 | 0.53 | **2.18** | **0.03** | **2.27** | **0.03** | 1.85 | 0.07 | 0.65 | 0.52 | 0.47 | 0.64 | 1.04 | 0.30 | -0.22 | 0.83 | -0.62 | 0.54 |
| Time point 3 | 0.01 | 0.99 | -0.03 | 0.97 | -0.15 | 0.88 | 0.13 | 0.90 | -1.03 | 0.31 | -0.13 | 0.90 | -0.41 | 0.69 | 0.27 | 0.79 | -1.25 | 0.22 |
| All time points | 0.25 | 0.80 | -0.02 | 0.98 | -0.17 | 0.87 | 0.19 | 0.85 | -0.33 | 0.74 | -0.12 | 0.91 | -0.43 | 0.67 | 0.32 | 0.75 | -0.46 | 0.65 |
| *COWAT: Letter fluency* | | | | | | | | | | | | | | | | | | |
| Time point 1 | -0.91 | 0.36 | **2.47** | **0.02** | **2.51** | **0.01** | **2.16** | **0.03** | 1.77 | 0.08 | 0.30 | 0.76 | 1.03 | 0.31 | -0.56 | 0.58 | 0.17 | 0.87 |
| Time point 2 | 0.94 | 0.35 | 0.81 | 0.42 | 0.78 | 0.44 | 0.74 | 0.46 | 1.41 | 0.16 | -0.36 | 0.72 | -0.07 | 0.94 | -0.52 | 0.60 | 0.78 | 0.44 |
| Time point 3 | 0.56 | 0.58 | 0.99 | 0.33 | 1.85 | 0.07 | 0.01 | 0.99 | 1.32 | 0.19 | -0.66 | 0.51 | 1.37 | 0.18 | **-2.52** | **0.01** | 0.51 | 0.61 |
| All time points | 0.07 | 0.95 | 1.75 | 0.08 | **2.19** | **0.03** | 1.05 | 0.29 | 1.93 | 0.06 | -0.11 | 0.92 | 1.12 | 0.26 | -1.22 | 0.24 | 0.77 | 0.44 |
| *COWAT: Category fluency* | | | | | | | | | | | | | | | | | | |
| Time point 1 | -1.51 | 0.14 | 1.11 | 0.27 | 0.77 | 0.22 | 1.40 | 0.17 | **2.15** | **0.03** | 0.14 | 0.89 | -0.15 | 0.88 | 0.40 | 0.69 | 1.93 | 0.06 |
| Time point 2 | 1.10 | 0.27 | 0.47 | 0.64 | 1.22 | 0.23 | -0.38 | 0.70 | 1.05 | 0.30 | -0.59 | 0.56 | 1.01 | 0.32 | -1.95 | 0.06 | 0.56 | 0.58 |
| Time point 3 | 0.91 | 0.37 | 1.10 | 0.28 | 1.64 | 0.11 | 0.41 | 0.68 | 1.84 | 0.07 | -0.57 | 0.57 | 1.02 | 0.31 | **-1.98** | **0.05** | 1.17 | 0.25 |
| All time points | 0.06 | 0.96 | 1.20 | 0.23 | 1.57 | 0.12 | 0.67 | 0.51 | **2.21** | **0.03** | -0.30 | 0.77 | 0.87 | 0.38 | -1.31 | 0.19 | 1.67 | 0.10 |
| *PCA 1* | | | | | | | | | | | | | | | | | | |
| Time point 1 | 1.08 | 0.28 | -1.34 | 0.18 | -1.24 | 0.21 | -1.36 | 0.18 | **-2.08** | **0.04** | 0.46 | 0.65 | 0.03 | 0.98 | 0.65 | 0.52 | -1.27 | 0.21 |
| Time point 2 | -1.02 | 0.31 | 0.56 | 0.58 | 0.16 | 0.87 | 0.91 | 0.37 | -1.02 | 0.31 | 0.72 | 0.48 | -0.02 | 0.99 | 1.12 | 0.27 | -1.33 | 0.19 |
| Time point 3 | 0.08 | 0.94 | **-2.10** | **0.04** | **-2.86** | **6.0 x $10^{-3}$** | -1.13 | 0.26 | **-3.80** | **3.6 x $10^{-4}$** | 0.08 | 0.94 | -0.02 | 0.99 | **2.13** | **0.04** | **-2.67** | **0.01** |
| All time points | -0.42 | 0.68 | -1.23 | 0.22 | -1.70 | 0.33 | -0.60 | 0.55 | **-2.23** | **0.03** | 0.68 | 0.50 | -0.64 | 0.52 | 1.48 | 0.14 | -1.35 | 0.18 |
| *PCA 2* | | | | | | | | | | | | | | | | | | |
| Time point 1 | -1.62 | 0.11 | 1.04 | 0.30 | 0.65 | 0.52 | 1.33 | 0.19 | 1.43 | 0.16 | 0.89 | 0.38 | 0.22 | 0.83 | 1.13 | 0.26 | 1.40 | 0.17 |
| Time point 2 | -1.79 | 0.08 | 0.59 | 0.59 | 0.14 | 0.89 | 0.99 | 0.33 | 0.40 | 0.69 | 1.13 | 0.27 | 0.26 | 0.80 | 1.52 | 0.14 | 0.62 | 0.54 |
| Time point 3 | -0.33 | 0.75 | 0.99 | 0.33 | 0.85 | 0.40 | 1.01 | 0.32 | 0.35 | 0.73 | -0.19 | 0.85 | -0.04 | 0.97 | -0.32 | 0.75 | -0.40 | 0.69 |
| All time points | -1.54 | 0.13 | 1.20 | 0.23 | 0.95 | 0.34 | 1.26 | 0.21 | 1.28 | 0.20 | 0.70 | 0.48 | 0.48 | 0.64 | 0.55 | 0.58 | 0.99 | 0.33 |

SDMT - Symbol Digital Modalities Test, PASAT - Paced Auditory Serial Addition Test, CVLT-II - California Verbal Learning Test-II, BVMT-R - Brief Visuospatial Memory Test Revised , CWIT - Color Word Interference Test , COWAT - Controlled Oral Word Association test, PCA - Principal Component Analysis
